# Cryptic diversity and impacts of domestication in the Black Soldier Fly (*Hermetia illucens*) genome

**DOI:** 10.1101/2023.10.21.563413

**Authors:** Tomas N. Generalovic, Christoph Sandrock, Benjamin J. Roberts, Joana I. Meier, Martin Hauser, Ian A. Warren, Miha Pipan, Richard Durbin, Chris D. Jiggins

**Affiliations:** Department of Zoology, University of Cambridge, Cambridge, UK; Department of Livestock Sciences, Research Institute of Organic Agriculture (FiBL), Frick, Switzerland; Georgina Mace Centre for the Living Planet, Faculty of Natural Sciences, Imperial College London, London, UK; Tree of Life Programme, Wellcome Sanger Institute, Wellcome Trust Genome Campus, Hinxton, Cambridge, UK; California Department of Food and Agriculture, Plant Pest Diagnostics Branch, Sacramento, CA, USA; Better Origin, Entomics Biosystems Limited, Cambridge, UK; Department of Genetics, University of Cambridge, Cambridge, UK

## Abstract

The Black Soldier Fly (*Hermetia illucens*) is the main species in the developing global industry of insects as food and feed, but little is known about its natural diversity or the genetic basis of its domestication. We obtained whole genome sequences for 54 individuals from both wild and captive populations. We identified two genetic lineages at least 3 million years divergent, revealing cryptic diversity within the species. Our study indicates that the most common populations used for commercial and academic applications are primarily derived from just one of these lineages, likely originating from a wild North American progenitor. We find that captive populations show strong reductions in genetic diversity, consistent with genome-wide effects of population bottlenecks and drift associated with rearing in captivity. Some limited evidence of gene flow between divergent lineages was observed, as well as evidence of hybridisation from domesticated populations into the wild. Our study suggests that natural genetic diversity could provide important variation for industrial purposes in this novel agricultural species.

**Author Summary:** Animals and plants have been domesticated by humans over thousands of years, shaping their genomes and yielding beneficial or improved traits for agriculture. Insect farming has recently reached industrial scales in order to recycle organic wastes into feed and food grade material. The Black Soldier Fly is the flagship insect species in this novel agriculture sector. However, the understanding of its evolutionary history and the genetic impact of intense farming has not been well studied to date. We examined whole genomes from populations sampled from across the worldwide distribution of Black Soldier Fly to investigate the impact of domestication and reveal the evolutionary history of the species. We found that farmed populations show distinct genomic regions showing reduced variation that could represent signatures of selection or demographic effects. We also identify two distinct lineages of Black Soldier Fly that have been diverging for millions of years, comparable to divergence between different species in other insect groups, yet show evidence of genetic exchange, indicating a lack of reproductive incompatibility in the populations tested. This work is the first to describe the genetic impact of domestication in the Black Soldier Fly and provide evolutionary insights using whole genomes. These results imply that considerable wild diversity could be harnessed for improving agricultural efficiency.

## Introduction

Domestication of plants and animals has enabled successful human population growth and globalisation [1]. Domestication is a coevolutionary process that emerges from a specialised mutualism where a species fitness is managed through manipulation of the environment, food-source, behaviour, and mate choice [2,3]. Evolutionary changes in physiological, morphological, and behavioural traits over time have led to specialised crops and animals with a host of phenotypic diversity, exemplified by *Brassica oleracea* cultivars and dog breeds [4,5]. This process of domestication also provides insights into evolution on short time scales [6,7]. Whilst domestication has been extensively studied in plants, mammals, birds and fish [4,5,8,9], the silkworm, *Bombyx mori*, is currently the only undisputed domesticated insect [10]. Here we focus on another candidate domesticated insect, the Black Soldier Fly, *Hermetia illucens* (L., 1758; Diptera: Stratiomyidae), in which both captive and wild populations can be found around the globe.

The Black Soldier Fly has only become an important and globally mass farmed species within the past two decades [11–13]. In response to rising food demands driven by an ever-increasing human population, novel and more sustainable agricultural practices are being developed, including the insects for food and feed industry [13,14]. The Black Soldier Fly can convert a diverse range of organic wastes into protein and fat rich biomass [11,15,16]. It has a cosmopolitan distribution with an origin in the Americas [17], most likely South America [18]. The global distribution of the Black Soldier Fly outside of the Americas is thought to be a result of human mediated dispersal and by 1960 the species had established most of its present-day distribution [17]. Microsatellite and mtDNA studies have shown that the species harbours high levels of genetic diversity [18–20], which suggests considerable potential for artificial selection and the development of improved strains [21,22]. However, commercial activity is currently dominated by a few closely related populations all derived from one strain founded in Georgia, USA in the 1990’s by Craig Sheppard [12,18]. Captive populations such as these have likely undergone or are currently experiencing a multitude of domestication bottlenecks and genetic adaptions associated with artificial insect farming conditions.

Whilst previous studies using microsatellites and mtDNA have provided initial insights into genetic diversity in Black Soldier Fly [18–20], these markers only sample a small proportion of the genome providing an incomplete view of genomic diversity [23]. In contrast, whole-genome sequencing captures variation genome-wide providing a comprehensive view of population diversity and structure [24]. Whole-genome sequencing is therefore able to provide more accurate estimates of genetic diversity, differentiation and patterns of linkage disequilibrium. Importantly, this enables genome wide patterns of drift and selection, signals that are typically undetectable with microsatellite data, to be detected. These features make whole-genome sequencing well-suited for revealing the evolutionary processes during domestication and within-species divergence.

Here we explore Black Soldier Fly diversity on a whole-genome scale, using a previously published high quality reference genome [25]. Using this first genome-wide dataset of the Black Soldier Fly, sequenced to high coverage, we explore the genomic landscape of captive and wild populations from key geographic locations. We address the following questions: (1) What are the phylogenetic relationships among Black Soldier Fly lineages? (2) When comparing genomes of captive populations to wild relatives can we reveal common patterns associated with domestication? (3) What are the genetic consequences of introgression in the wild and within insect mass-rearing commercial facilities?

## Results

### Genome sequencing and variant calling

We sequenced 54 Black Soldier Fly genomes to an average 22x coverage alongside an additional outgroup species, *Ptecticus aurifer* (supplementary table 1). To capture the extent of genetic variation within the species we sampled Black Soldier Fly individuals from genetically distinct (captive and wild) populations, previously identified from microsatellite data [18]. The remaining dataset consisted of representative samples of active industrial and academic captive lines. In addition, we sampled wild sourced populations as part of a taxonomic voucher set including a single outgroup species. We hereafter term academic/industry-derived samples as ‘captive’ (*n*=35/54) whereas samples directly sampled from in the field were termed ‘wild’ (*n*=19/54) (supplementary table 1). Captive individuals belonging to South America, Africa and Australia were recorded as wild caught and recently established as a captive colony. After variant calling and stringent quality filtering, we identified 33,940,069 bi-allelic Single Nucleotide Polymorphisms (SNPs).

### Divergent lineages and cryptic species complex

Phylogenetic analysis on the genome-wide SNP data showed two major genetic lineages. The first lineage contains wild sampled individuals from North America and many captive populations, hereafter named lineage-*a* (Figure 1A). The captive populations found in lineage-*a* are made polyphyletic by the North American wild population. European and North American captive populations form sister clades in line with a documented split from an original founding population, named hereafter as the ‘Sheppard’ strain. The ‘EVE’ genome reference line also appears to be a derivative of the Sheppard strain. The second lineage, lineage-*b,* is formed from a diverse range of wild and captive populations. In lineage-*b*, wild and captive clades broadly cluster according to geographic location, indicating these captive populations are established from locally occurring populations, as documented. Individuals from Asia (wild) and Australia (captive) also form an ’Australasian’ clade. Phylogenetic methods using coalescence-based methods (ASTRAL) with input trees calculated using either Neighbour-Joining (NJ) or Maximum-Likelihood (ML) algorithms also support a split between major lineages (supplementary figure 1A & 1B). However, the NJ approach suggests lineage-*b* samples are paraphyletic with respect to lineage-*a* individuals (supplementary figure 1A).

**Figure 1.**
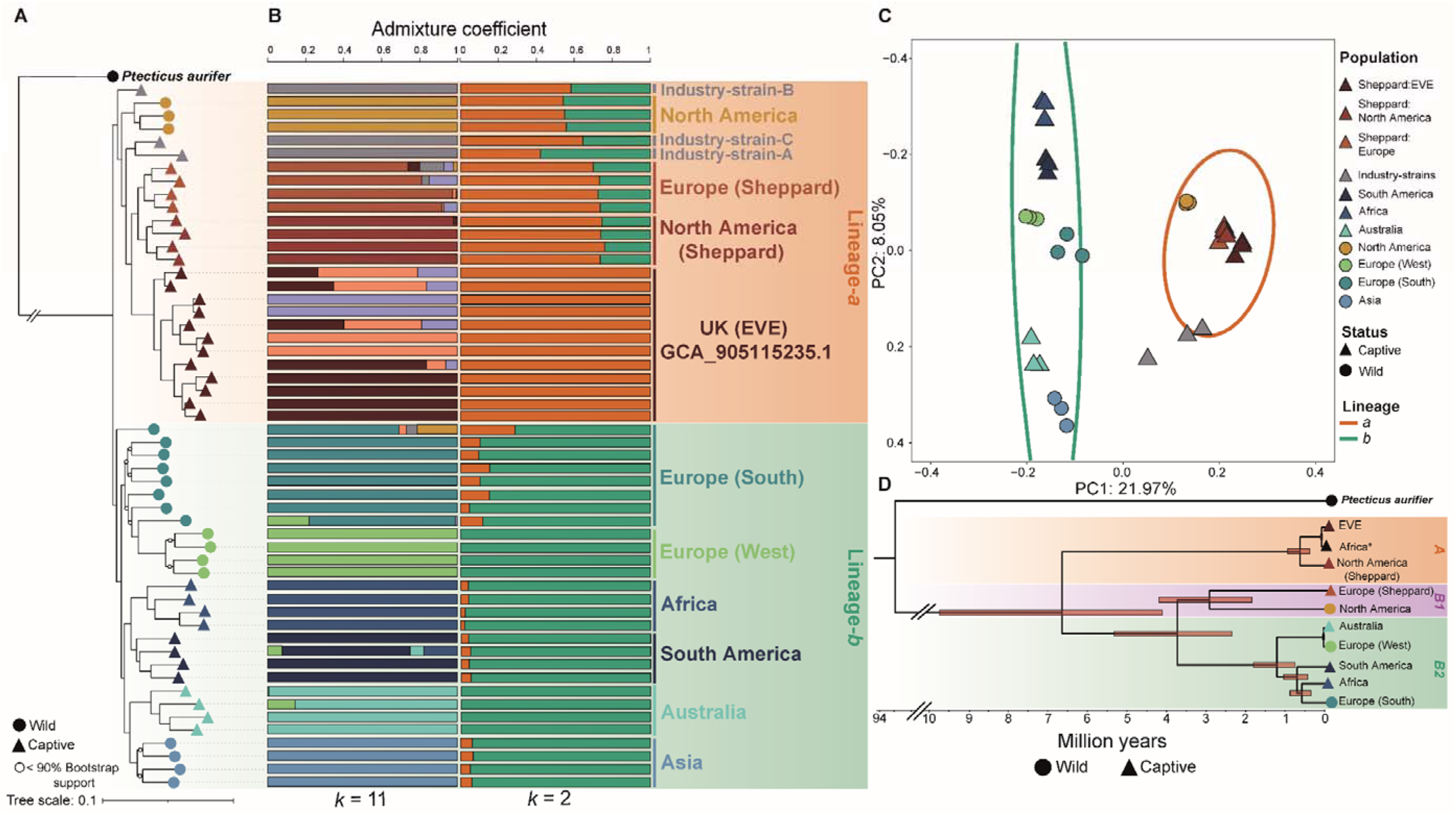
(**A**) Maximum Likelihood phylogenetic reconstruction using whole Black Soldier Fly (n=54) and *Ptecticus aurifer* (n=1) genomes. (**B**) ADMIXTURE analysis of the Black Soldier Fly population structure at *k*=11 and *k*=2 (lowest cross-validation error support) indicating pronounced population structure. (**C**) Principal Component Analysis (PCA) using a subsampled dataset of whole genomes (n=3 per population) supporting two discrete genetic lineages. A 95% confidence interval (CI) ellipse is drawn over major lineages based on phylogenetic (**A**) and clustering (**B**; *k*=2) assignment of individuals. (**D**) BEAST time divergence model representation of the Black Soldier Fly mitochondrial phylogeny reconstructed using a single whole mitochondrial (mt) genome from each major clade identified in the whole-genome phylogeny (**A**). Lineages recovered in the mtDNA tree are labelled including the unresolved mtDNA lineage-*B*. *Single mt genome obtained from [20].

Evidence for substantial divergence between the two nuclear lineages was also supported by ADMIXTURE (Figure 1B) and Principal Component Analysis (PCA; Figure 1C). Optimal clustering in ADMIXTURE analysis was identified at *k* = 2 (Figure 1B; supplementary table 2), and this was consistent when subsampling an equal number of individuals per population (supplementary figure 2). A similar Cross Validation (CV) rate was identified at *k*=3 (0.887) to *k*=2 (0.885) indicating a third clustering group primarily consisting of EVE and captive North American individuals (supplementary figure 3). At higher *k*-values, *k* = 11 (Figure 1B), wild European populations show distinct clusters indicating a diverse within continent population structure consistent with independent colonisation events and likely subsequent patterns of isolation-by-distance. A similar pattern is observed across a larger geographic range in Asian and Australian samples in agreement with the successive colonisation of Australasia [18,20]. PCA analysis also showed two main clusters with many captive populations clustering with wild North American samples while the remaining wild and captive samples found from a broad geographic range form a second major cluster (Figure 1C). Several individuals belonging to various industry populations (Ind-A, -B & -C) are found in the PCA as intermediates between lineages, suggesting admixed origins for these populations. When grouped by discrete populations identified in clustering analysis we found moderate genome-wide levels of *F_ST_* (0.178) and *d_XY_* (0.081) between nuclear lineages. This suggests a long history of evolutionary separation with limited gene flow between lineages. Within lineages, lineage-*b* had higher genome-wide values of both *F_ST_* and *d_XY_* as compared to lineage-*a* (supplementary figure 4).

### Ancient divergence between Black Soldier Fly lineages

We produced *de novo* mitochondrial assemblies for the 54 Black Soldier Fly individuals and one *P. aurifer* individual (supplementary table 3; supplementary figure 5). The *de novo* Black Soldier Fly and *P. aurifer* assemblies are comparable to published reference mitochondrial genomes (supplementary table 3; supplementary figure 6) [20,26,27]. In contrast to the two lineages (*a* & *b*) identified from nuclear sequences, three major lineages were recovered in the mtDNA data (*A*, *B1* & *B2*) (Figure 1D & supplementary figure 6). Nuclear lineage-*a* samples fell into mitochondria clades *A* and *B1*, in which lineage-*A* diverged an estimated 4.1 to 9.7 million years ago (my; 95% Confidence Intervals (CI); mean 6.8 my) (supplementary table 4). Lineage-*B1* of the mtDNA tree contains nuclear lineage-*a* wild North American and captive European-Sheppard populations (Figure 1D). mtDNA lineage-*B2* contains all samples from nuclear lineage-*b* and diverged from lineage-*B1* approximately 2.4 – 5.3 my ago (95% CI; mean 3.8 my) (supplementary table 4). Confidence in the models dating outputs is high, in the initial calibrating dating run, comprising a wider dipteran phylogeny, all calibrated nodes fit well within the assigned constraints whilst non-calibrated nodes show agreement with existing literature (supplementary table 4), suggesting that the dating model has performed well. Here, we dated the Stratiomyidae - Tabanidae split at 182.0 my ago and the Stratiomyidae root at 81.1 my whilst these splits have been previously dated at around ∼190 my and ∼75 my ago respectively (see supplementary table 4 and references within), well within the 95% confidence intervals. We also generated Neighbour-Joining (NJ) trees of Cytochrome Oxidase I (COI) markers, such that additional previously published samples could be incorporated, including some from wild Central American populations (supplementary figure 7). This confirms that Sheppard-derived populations of lineage-*a* are most closely related to wild samples from North America.

To corroborate the divergence times between lineages identified in the mtDNA data we performed additional dating analysis using whole genome evidence. Using the known split, between *P. aurifer* and *H. illucens,* we constrained the phylogeny to predict the divergence between nuclear lineages-*a* and -*b*. Our whole-genome analysis predicts that the major split between lineages was estimated between 3.9 – 13.8 my ago (95% CI; mean 8.9 my) (supplementary figure 8). Whilst the mean whole-genome divergence between lineages is higher, in both mtDNA analysis provided above and in published work [20], the majority of the range (3.9 -13.8 my) overlaps considerably with the mtDNA confidence intervals of 4.1 - 9.7 my and further suggests considerable diversity in the Black Soldier Fly.

### Genetic signals of domestication

Genomic signals of domestication are typically characterised by genome-wide reductions in genetic diversity and elevated linkage disequilibrium (LD) associated with a domestication bottleneck and small effective population sizes (*Ne*). On average, genome-wide heterozygosity of captive Sheppard (0.031) strains shows a 39% reduction compared to wild populations (0.051), supporting the domestication hypothesis (supplementary table 5; supplementary figure 9). Additional captive populations from South America exhibit the highest levels of diversity (0.087), as expected if this represents the ancestral range of the species despite captivity status, followed by African populations. In general, extensively reared captive populations showed lower diversity with the exception of the European population of the Sheppard strain which showed levels of diversity similar to that of wild populations. Within-population estimates of LD revealed elevated levels in captive populations consistent with a genetic bottleneck during domestication (supplementary figure 10). This analysis highlighted genome-wide patterns of domestication in captive populations.

Due to novel selection pressures and somewhat recent bottlenecking during domestication, genomes of captive populations would be expected to contain higher frequencies of homozygous genomic regions, due to a loss of rare alleles, compared to the wild. We generated evidence for genomic changes associated with domestication and demographic events by identifying regions of reduced diversity by calculating Runs of Homozygosity (RoH) (supplementary table 6 & 7). In total, captive populations (*n* = 5,245) contained a higher number of RoH than wild (*n* = 352) based populations. However, to quantify unique RoH we reduced overlapping regions into single counts that were present in all individuals within captivity status groups. We identified a total of 907 captive and 53 wild unique RoH with 245 shared RoH regions (supplementary figure 11). Although only 53 RoH regions were unique to wild populations, these accounted for just 6.5 Mbp, whilst 80.9 Mbp (83%) of the wild RoH regions were shared with captive populations whereas captive populations span 623.0 Mbp of the genome. Captive RoH were significantly more numerous (Wilcoxon rank-sum test, *W* = 445, *p* < 0.001) and on average longer (Wilcoxon rank-sum test, *W* = 444, *p* < 0.001) at 0.69 Mb, indicating recent bottlenecks, compared to both shared (0.33 Mb) and wild (0.12 Mb) populations. We also performed a permutation test based on 10,000 randomisations across the genome to assess if the total unique captive RoH regions had more or less coverage than expected by random across the genome. Interestingly by assessing the unique captive RoH regions we found significantly lower than expected (*p* < 0.001, *Z* = -45.4). Whilst numerous and long, this result suggests that captive RoH’s appear tightly clustered and less dispersed across the genome indicating hotspots impacted by demographic or selection processes (supplementary table 8). However, many of these patterns are somewhat driven by extensive RoH within the EVE population. We identified six regions in which RoH clustered consistently in at least 45% of the captive individual samples (Figure 2).

**Figure 2.**
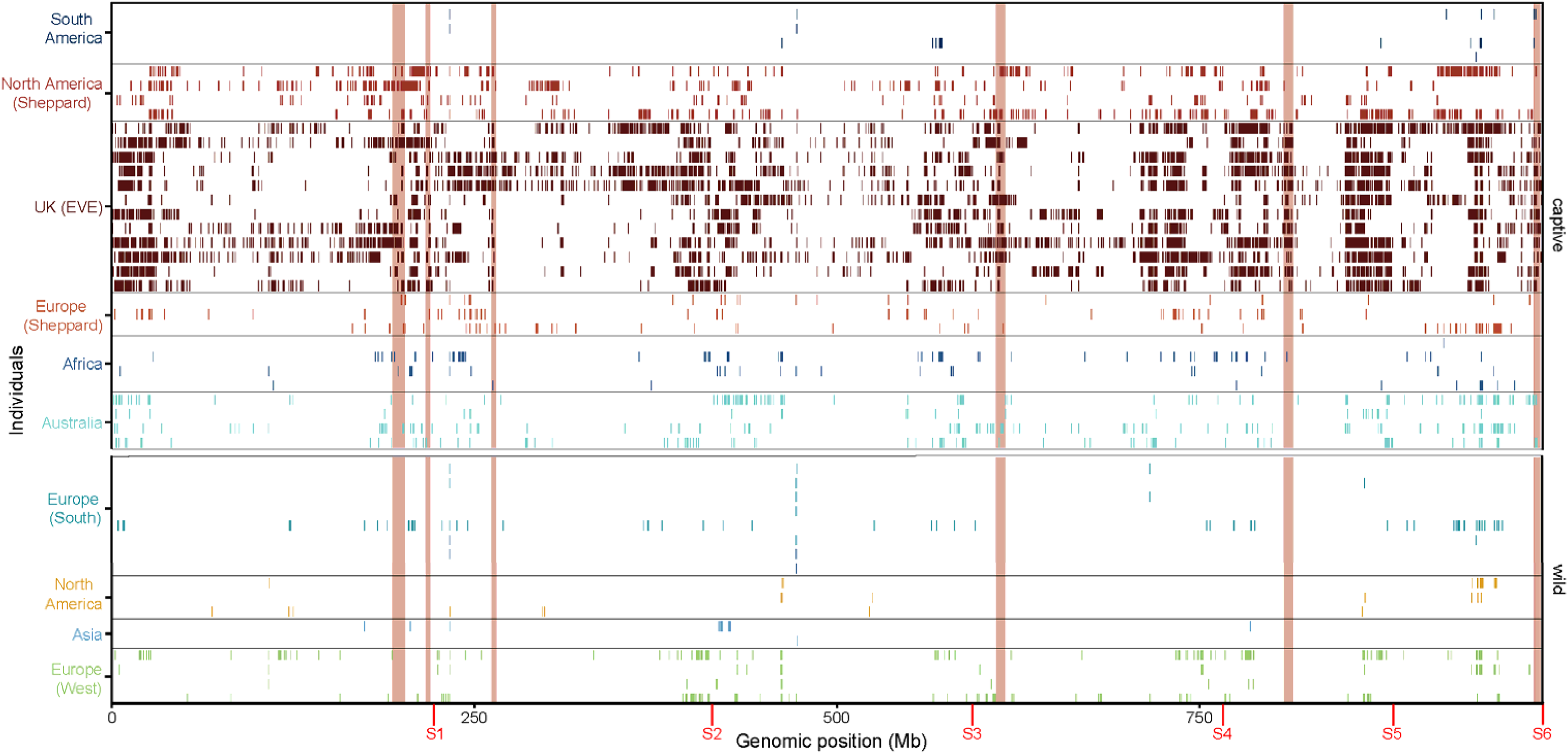
Runs of Homozygosity calculated across the genomes of wild and captive Black Soldier Fly populations. Each row in the y-axis represents an individual and colour represents the population. Chromosome boundaries are provided in red on the x-axis along with genome position (Mb). Regions clustered with RoH are highlighted in coloured blocks highlighting regions in which RoH cluster in over 45% of the captive individuals on chromosomes S1 (*n*=2), S2 (*n*=1), S4 (*n*=1), S5 (*n*=1) and S6 (*n*=1).

We also used genome-wide scans of genetic diversity (lf) and differentiation (*F_ST_*) to further investigate the genomes for differences between captive Sheppard populations and their closest sampled wild relative, North America. Strong patterns of genome-wide nucleotide diversity between captive and wild populations were relatively absent with North America (wild) and South America (wild) both showing higher levels of diversity whereas captive populations were reduced (supplementary figure 12). Elevated *F_ST_* was observed genome-wide for all three populations, however, strong peaks were observed converging on chromosome five for all three populations (Figure 3). Notably, there was strong *F_ST_* peaks overlapping in the Sheppard North American and European population on chromosomes one, four and five. The EVE population showed broad and elevated *F_ST_* peaks, for example, the large RoH locus on chromosome five previously identified in the EVE population had broadly elevated *F_ST_*but was not identified in other Sheppard populations suggesting this locus is likely demographic in nature and specific to the EVE population. Combined, these data suggest that domestication has left a strong signature on the genomes of captive populations, either through drift or selection.

**Figure 3.**
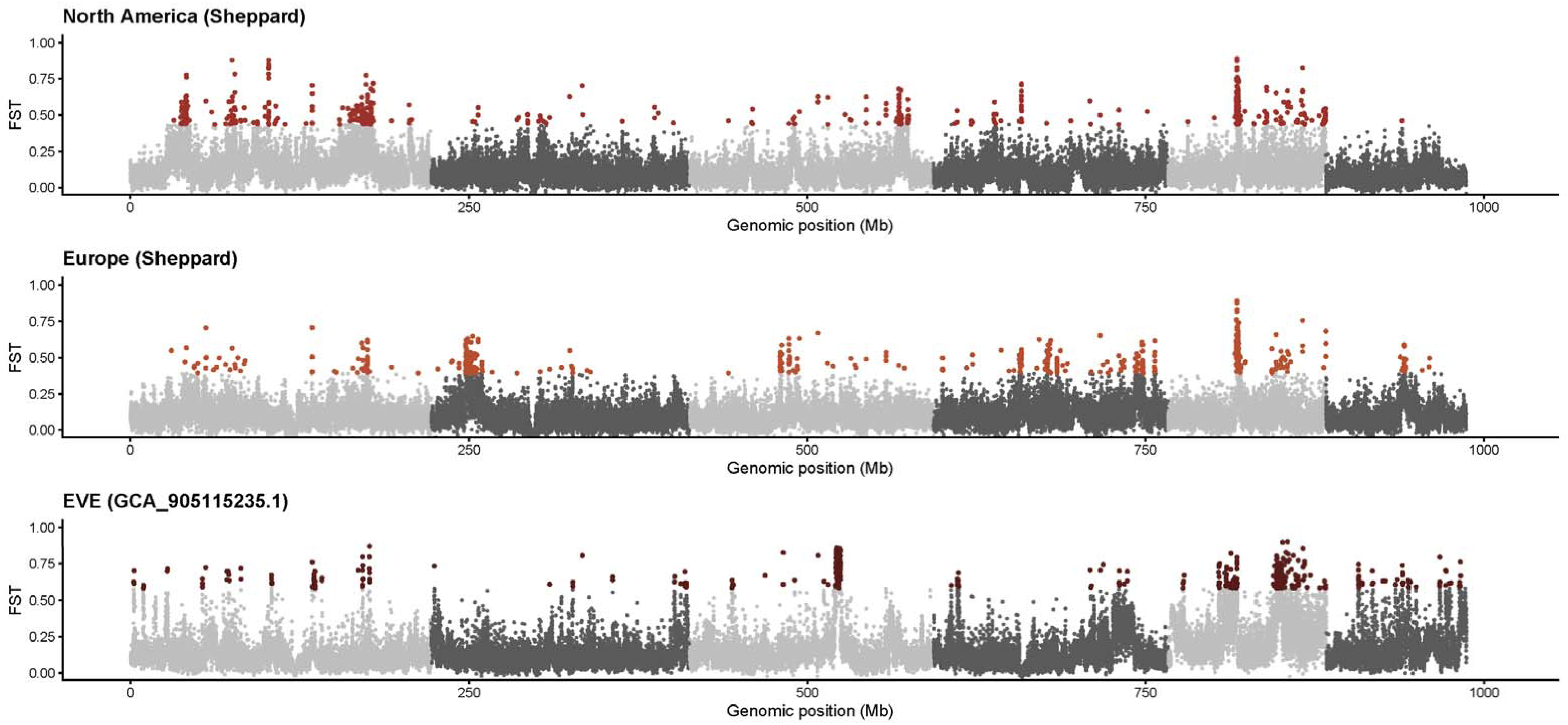
Genetic differentiation (*F_ST_*) between various domesticated Black Soldier Fly populations. Sheppard strain from North America, Europe, and the EVE populations are compared to the closest wild sampled progenitor population from North America. Outlier regions identified within the top 99^th^ percentile of 20 kb windows containing SNPs are indicated using population specific-colored points.

### Genomic consequences of introgression

Evidence for introgression between the major lineages of Black Soldier Fly comes from a comparison between nuclear (n=54) and mitochondria (n=112) whole-genome phylogenies, which showed mito-nuclear discordance (Figure 4). We found 12 samples with mito-nuclear discordance at the broad level of the major lineages and clades discussed above; all but two of these were localised to mtDNA clade-*B1*. However, overall tree discordance was low affecting only 22% of sampled individuals.

**Figure 4.**
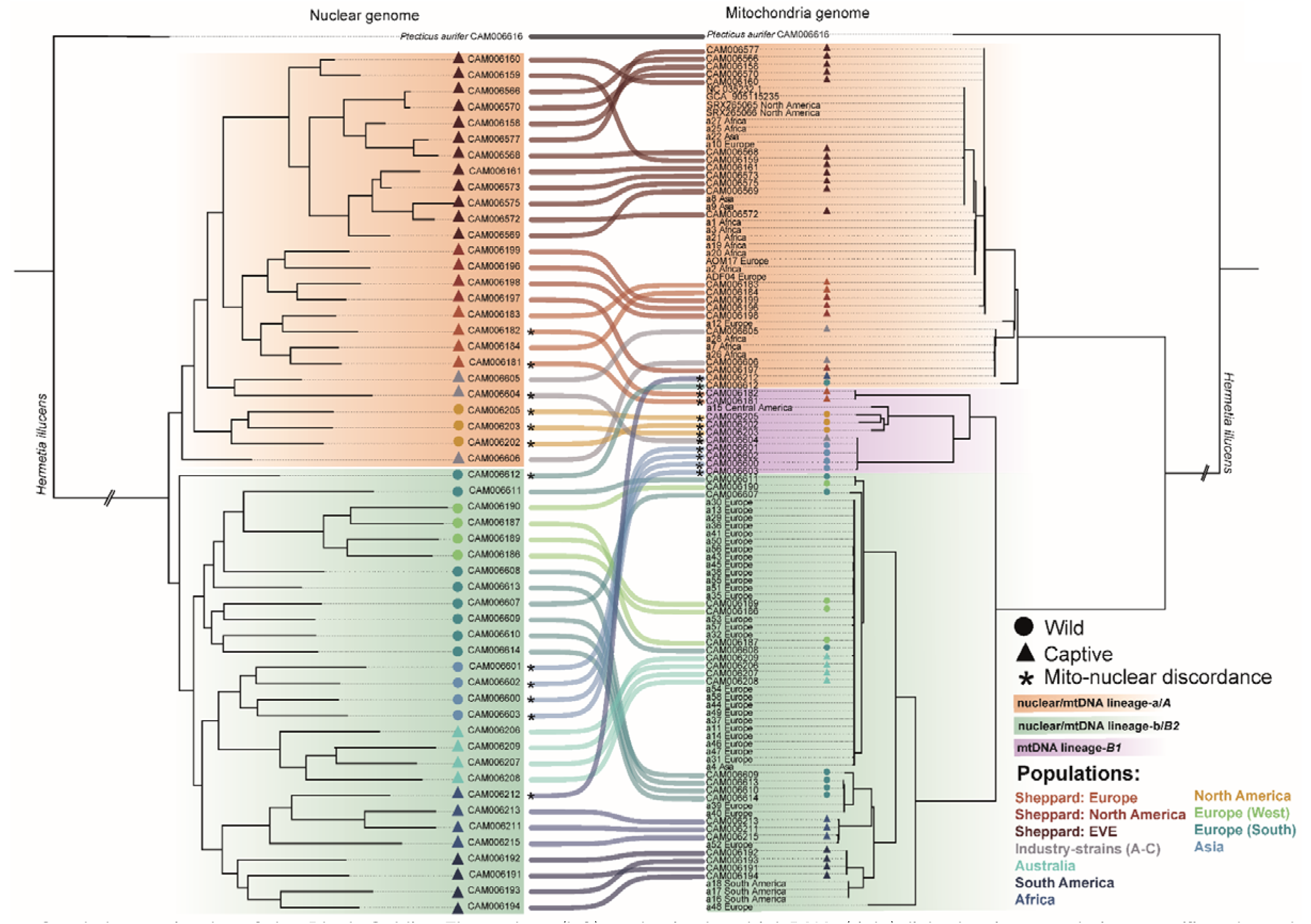
Co-phylogenetic plot of the Black Soldier Fly nuclear (left) and mitochondrial DNA (right) linked using population specific-coloured ribbons and individual CAM ID’s. Evidence of mito-nuclear discordance is described by an asterisk in individuals that cross the major identified lineages (*A*, *B1* & *B2*). Whole mitochondria genomes from this study (**I**) and various (**II**) public repositories are used to expand the mtDNA tree.

We next examined genome-wide patterns of introgression within and between divergent lineages. We generated *f*-branch (*f_b_*) statistics using the whole-genome phylogeny of the Black Soldier Fly to estimate genome-wide allele sharing patterns between major lineages using *P. aurifer* as the distant outgroup. Generally, there was little evidence for gene flow between the major lineages, with a few notable exceptions (Figure 5A).

**Figure 5.**
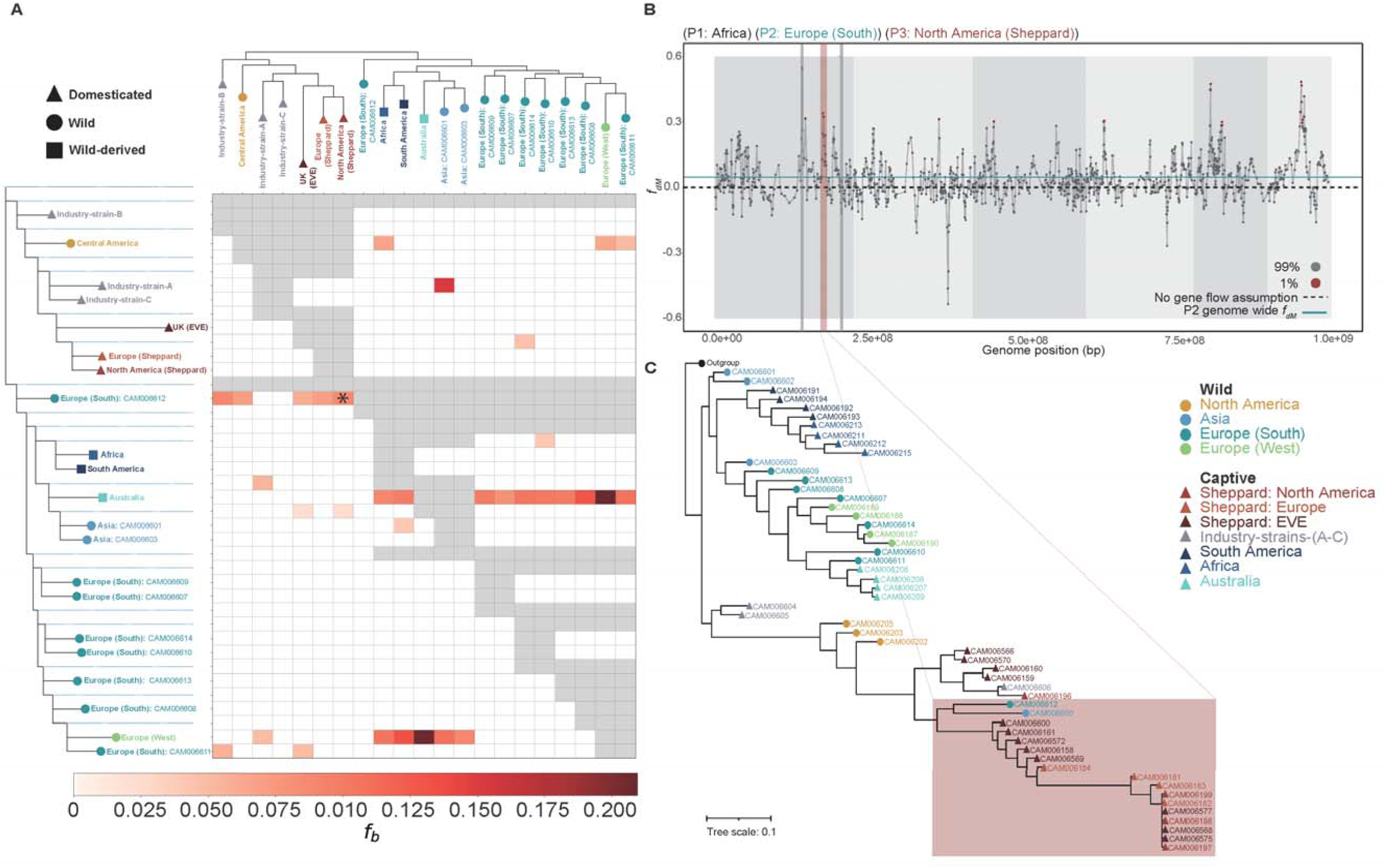
(**A**) Admixture analysis using the *f*-branch (*fb*) statistic across sampled Black Soldier Fly individuals indicating excess allele sharing as indicated by utilised species trees (left and top). Phylogenetic tree on the y-axis represents populations P1 & P2 and is expanded to indicate internal branches enabling comparisons to the P3 population along the x-axis. Admixture is detected between P2 & P3 with P1 fixed as the closest sister group. Grid cell colour and intensity indicates levels of significant allele sharing. Grey shading indicates where the tests cannot be performed due to tree topology. Significant values only are plotted. *Indicates zoomed example for panels B & C. (**B**) Genome-wide *fDM* plotted highlighting 1% outlier values of allele sharing between P3 and P2. Alternating light and dark grey shading indicate autosomes. (**C**) Local gene tree of the introgressed region (2.1 Mb; grey) from the North American Sheppard Black Soldier Fly population (P3) into the Southern Europe wild (P2) individual showing clustering with captive populations (highlighted block).

The strongest evidence of introgression from lineage-*a* into *-b* (*f_b_* = 0.06 - 0.09) occurred from captive lineage-*a* into a wild southern European (CAM006612) representative (Figure 5A). This same individual was previously identified as showing mito-nuclear discordance (Figure 4). This may reflect introgression due to captive strains escaping into the wild. To identify specific genomic regions implicated, we used *f_dM_* to scan the genome, revealing genome wide introgression signals peaking over chromosome one (Figure 5B). Of the introgressed genetic material into the wild, 93% of the genes in this 2.1 Mb region were identified as Cytochrome P450s (supplementary table 9). Gene trees group introgressed wild individuals within a Sheppard-derived captive clade, corroborating the hypothesis of introgression (Figure 5C). Within lineages, we found evidence for introgression hotspots between lineage-*b*’s wild Europe and captive Australia individuals from multiple widespread locations, likely a product of anthropogenic mediated dispersal (Figure 5A). We also found evidence for bi-directional gene flow between wild western Europe and Australia indicative of repeated exchange between these geographically distant locations. This evidence supports similar observations of Black Soldier Fly colonisation of western Europe via admixed populations including Australasian ancestors [18].

We also detected introgression (*f_b_* = 0.152) into industry-strain-A (Ind-A) from wild Asian populations (Figure 5A). Genome-wide scans of *f_dM_* in Ind-A suggests a primarily lineage-*a* genetic background, indicated with a negative genome-wide mean *f_dM_*, with a large 7.07 Mb region (S3: 150867943-157941741) of wild Asian origin introgressed on chromosome three (supplementary figure 13), containing a total of 121 genes (supplementary table 10). Several genes across this region have multiple copies including Collagen alpha (*Col4a1*; n=2), Dual oxidase (*Duox*; n=2) and salivary secreted peptide (*Sgsf*; n=5) genes (supplementary table 10).

## Discussion

Whole-genome sequencing of previously identified genetic clusters from populations of the Black Soldier Fly (*Hermetia illucens*) has provided insight into the natural diversity of this agriculturally important species. We have identified patterns associated with the genomic footprint of domestication. We find domesticated populations of lineage-*a* are largely derived from a North American wild population whereas lineage-*b* captive individuals appear to be naturalised within geographic regions likely due to their recent captivity or ongoing gene flow with the wild. Genetic diversity within this species is high and there are two highly divergent nuclear lineages. One of these lineages (-*b*) presents significant opportunity for establishing breeding programs due to its high genetic diversity in contrast to lineage-*a*, the lineage currently in widespread use by industry. Despite a worldwide distribution we observe an overall lack of hybridisation between these diverse lineages apart from a few examples of localised gene flow. Nonetheless, this may present opportunity for breeding strategies introducing novel wild genetic diversity into captive populations to improve performance [28]. The deep divergence between lineages may pose challenges for interbreeding programs due to hybrid breakdown, reduced fitness or pre-mating isolation, although recent crossing experiments suggest little evidence for incompatibilities [29].

### Revealing cryptic genetic diversity

Whole-genome analysis reveals deep structure within the Black Soldier Fly. Previous mtDNA evidence estimated divergence between major clades over 2 my ago [20], but these data indicate even deeper divergences up to 6.8 – 8.4 my ago, greater than the divergence between *Drosophila* species [30–37]. Combined with a lack of evidence for extensive admixture, our data would suggest that Black Soldier Fly lineages could represent distinct species. Nonetheless, the genetic evidence indicates ongoing hybridisation suggesting a lack of complete reproductive incompatibilities [18,19], consistent with results from laboratory crosses [29]. Further behavioural and morphological analysis would be required before these lineages can be formally described as species or breed logs can be defined.

The existence of cryptic lineages could have important implications for the industry [38–40]. Progress in breeding programs could be hindered if incompatibilities and/or pre-mating isolation exist between strains [41,42]. Cryptic lineages within the Black Soldier Fly are likely in part responsible for the high phenotypic variance previously described [43], including nutritional [44–46] and performance variation [47,48]. Previous work has demonstrated significant phenotypic variation among strains of Sheppard and Australasian origin [49]. Within lineage-*a,* different populations and their interbreeding likely also contributed to this phenotypic variation reported across various studies. Among these samples, lineage-*b* is more diverse and might therefore hold a comparatively larger pool of unrecognised phenotypic variation of possible commercial relevance. Whereas at present most industrial activity harnesses lineage-*a* [18], emerging evidence suggests that identifying and optimising strain-substrate combinations could significantly improve performance, highlighting the importance of further research into evolutionary divergence and phenotypic plasticity of this species in the case of industrial application [49,50]. Overall, it remains unclear the extent to which commercially motivated worldwide exchange will contribute to the breakdown of these divergent lineages. However, deciphering the evolutionary history of the Black Soldier Fly will contribute to improving breeding and targeted genomic selection programs.

Our data also provides insights into the biogeographic history of this species. Whilst the within-species lineage divergence time is suggested to be older in the whole-genome data (3.9 – 13.8 my) unknown mutation rates and demographic history, including the differences in generation time between domesticated and wild populations may have inflated this prediction. Nonetheless, both mtDNA and whole-genome data in this and previous studies all show deep divergence within the Black Soldier Fly [20]. The deep divergences observed (>3 my ago) predate recent anthropogenic expansion, suggesting that this must represent divergence originating in the original native range in the Americas [17,18,20]. This could be a result of isolation-by-distance (IBD) across a broad range from South to North America. Alternatively, deep divergence in the mtDNA and whole genome trees might reflect biogeographic events within the Americas. For example, divergence in mtDNA that occurred during the late Miocene period (∼6 mya) coincides with uplift of the Andean Plateau, potentially driving reproductive barriers between South American populations [19,51–53]. The split between mtDNA lineages-*B1* and -*B2* occurred more recently (∼3.2 my) which may represent colonisation of Central America after the rise of the isthmus of Panama [54]. Alternatively, the pattern of mito-nuclear discordance would be consistent with an introgression event over 3 million years ago whereby mtDNA lineage--*B1* introgressed into nuclear lineage-*a* [55]. Ancient hybridisation has been shown to facilitate adaptive ecological radiation and speciation which may have played a role in the diversification of this species [56]. Likewise, hybridisation may have played an important role in shaping the genetic basis for domestication and consequently, an entire industry largely building on lineage-*a* descendants. *Hermetia* are distributed throughout the globe, yet ∼65% of species are found in the Neotropics [57], and only the Black Soldier Fly (*H. illucens*) is commonly studied. Therefore, *Hermetia* species are an interesting system to explore the consequences of hybridisation on geographic and ecological diversification. However, divergence time estimates should be interpreted with caution due to the absence of a Black Soldier Fly mutation rate. The mutation rate was inferred from *Drosophila*, which split from *Hermetia* ∼200 mya. Variation in generation time and species-specific molecular mechanisms may have introduced significant error in these approximations [58].

### Genetic impacts of domestication

Black Soldier Fly domestication has led to a loss of genetic diversity (39%) approaching that of domesticated crops (47-72%) [59,60]. This is perhaps surprising considering the recent timescale, estimated between one and two decades, of Black Soldier Fly domestication. Continued domestication will likely exacerbate this reduction of genetic diversity in commercial facilities raising further concerns around inbreeding and disease susceptibility [61,62]. Our whole-genome nuclear analysis identifies North America as the likely progenitor of the common domesticated Black Soldier Fly Sheppard strain. Using published COI markers combined with this data further supports a progenitor of domesticated Black Soldier Fly to a wild North American origin [18–20]. This likely represents the single domestication event previously documented which established a colony over 200 generations ago [18]. In addition, we find evidence for at least a second independent domestication event, maybe more, occurring in captive populations in South America, Africa and Australia. Considerable genetic diversity and varying signals of domestication are present within described strains, potentially indicative of specific breeds. Whilst characteristic traits associated with distinct strains/ breeds have begun to be revealed, they remain largely unknown [43,49,50].

Unsurprisingly, over two decades of domestication occurring in parallel across the globe has shaped the genomes of farmed populations. Some caution should be applied when interpreting between signals of selection or drift, particularly when regions stretching over extreme distances are found, as inbreeding can also lead to runs of homozygosity and signatures similar to selective sweeps. Furthermore, our sample size for all populations apart from EVE were too small to reliably detect selective sweeps and therefore was not performed. Therefore, we analysed patterns of homozygosity associated with inbreeding, drift and selection and identified regions indicative of non-random genomic processes at play. Nonetheless, evidence suggests genomes of captive populations have been shaped by domestication and at least some of these regions may be signatures of selection. Although we cannot fully disentangle the contributions of selection and drift further research is required to disentangle these signatures with greater sample sizes. Despite this, domestication has played a significant role in shaping the genomic landscape of the Black Soldier Fly.

Domestication of the Black Soldier Fly may not only drive positive selection but, in the presence of abundant feed availability and optimal rearing conditions, may also lead to relaxed selection. Due to the nature of the industry, in which core breeding colonies are typically reared on nutrient rich substrates, it is possible that domesticated strains could lose the ability to effectively feed on nutrient poor organic waste substrates, threatening efficient Feed Conversion Ratios (FCRs). This may reposition these lines useful for insect waste management over nutrient upcycling. On the other hand, increased rearing densities for commercial efficiency may also drive positive selection for stress tolerance and disease susceptibility [63]. The use of whole-genome sequencing data has provided high resolution insights into the impact of domestication on the genome of the Black Soldier Fly, previously undetected by microsatellites and mtDNA. Although it is challenging to differentiate drift and selection, it is important to further identify candidate genomic regions that could be explored further to understand domestication.

### Evidence of introgression and genomic consequences of gene flow

Overall, there was limited evidence for introgression between major lineages of the Black Soldier Fly. In part, this may represent a bias in sampling as individuals representing major genetic clusters in the microsatellite data were chosen for sequencing here [18]. Despite this sampling, evidence of hybridisation was identified between lineages occurring in a small number of individuals in Europe, likely as a product of escaped flies from farms [20]. Genetic material found introgressed into a wild southern European individual on chromosome one showed an overrepresentation of P450 genes. This locus perhaps has an adaptive benefit in response to high mycotoxin concentration exposure in industrial organic waste streams [64,65]. However, these introgressed regions must undergo phenotypic validation to confirm their adaptive role. Whilst a lack of biosecurity measures allowing the release of domesticated flies into the wild poses conservation concerns, it also threatens potential undocumented and economically relevant trait loss from naturally diverse wild populations through genomic homogenisation [66,67]. In addition to genetic concerns, such uncontrolled releases may have broader ecological consequences. Domesticated populations may harbour unique disease threats to wild locally adapted populations [68].

There was also some limited evidence for gene flow into active commercial lines belonging to lineage-*a* from wild populations. Wild populations are notoriously difficult to establish in the lab but might harbour valuable phenotypic diversity that could be harnessed by developing introgressed lines through controlled breeding [61,69]. For example, we identified an introgressed genomic region derived from a wild Asian source within one industry population. All industry strains (A, B & C), sourced from Asian suppliers, showed evidence for admixture between lineage-*a* derived domesticated strains and wild Asian populations. Previous work has shown that admixed lineages are prevalent in production facilities, particularly across Asia [18]. Within an industrial setting introgression may promote genetic diversity [70] and enable the introduction of specific genes for novel applications [71]. Whilst extensive introgression may protect against colony collapse in the short term, establishing an industry with a single founding stock population with a highly similar genetic background severely increases the risk of disease susceptibility in the long term [61,62,72,73]. Clearly further work is needed to understand the degree of heterosis versus hybrid breakdown in crosses between divergent lineages [28]. In the light of this genetic diversity, cost effective and accessible genetic management will likely play an important role in this industry in the future.

### Conclusions

Our study indicates the existence of previously undescribed cryptic diversity within the Black Soldier Fly. Levels of divergence observed are in the same order of magnitude as divergent *Drosophila* species. This suggests a potential need for a taxonomic revaluation or breed book record keeping of the Black Soldier Fly following further sampling efforts in the Americas. We demonstrate that two decades of extensive captive rearing, originating from a probable wild North American progenitor, has led to dramatic changes in the genome. Importantly, irrespective of their distribution, captive rearing has left distinct signatures associated with domestication across genetically distinct populations. This species is an excellent system to study evolution due to the rapid and ongoing domestication pressures on an industrial and global scale. The genomic diversity described here sets the stage for future research to better characterise the genetic identity of strains and to monitor inbreeding in an industrial setting. Further sampling will be needed to elucidate the genetic basis of domestication, but this species could become an interesting case study for understanding human impacts on domesticated species as well as providing a foundation for genetic improvements.

## Materials and Methods

### Sample collection, whole genome sequencing and variant calling

We obtained 54 whole-bodied (adult) Black Soldier Fly and one *Ptecticus aurifer* sample for whole genome sequencing (WGS). Individuals were obtained from either previously sampled collections [18], maintained (but sourced from three independent Asian suppliers) within industry conditions (Better Origin, UK) or from a taxonomic voucher set (source: M. Hauser). Total genomic DNA was extracted from thorax muscle tissue and purified using the Qiagen Blood & Culture DNA Midi Kit (Qiagen, Germany). Libraries (*n* = 39) were prepared, and genome sequenced by BGI (BGI, Hong Kong) as in Generalovic et al. (2021), including the 12 previously published genomes. Novogene (Cambridge, UK) constructed a second set of libraries (*n* = 11) and performed paired end (PE) 150 bp sequencing on the NovaSeq platform. The remaining five libraries were sequenced in the same manner but constructed using a custom Tn5 based protocol [74]. Raw sequencing data was quality checked, with FastQC v0.11.4 [75] and visualised in MultiQC v1.9 [76] both pre- and post-adapter trimming with cutadapt v1.8.1 [77]. Treated reads were mapped to the GCA_905115235.1 reference genome [25] using bwa-mem v0.7.17 algorithm [78]. Resulting BAM files were sorted, duplicates marked and removed using samtools v1.9 [79] and picard v2.9.2 (http://broadinstitute.github.io/picard). Variant calling was performed following the best practices of Genome Analysis Toolkit (GATK) v3.7.0 [80]. Genotyped SNPs were quality checked and filtered for high quality (>75) sites setting a minimum and maximum depth cut-off (10-34X) and ignoring sites containing a high proportion of missingness (>15%). Assessment of contaminants based on allelic balance within samples was carried out using allelicBalance.py (https://github.com/joanam/scripts/blob/master/allelicBalance.py). An additional VCF dataset was generated using a minor allele frequency (MAF) filter threshold of 0.1.

### Mitochondrial *de novo* assembly

Analysis of mitochondrial genomes was performed by assembling *de novo* whole mitochondrial genomes using a subset of the novel WGS reads produced in this study and collected from previous work [20,25,26]. We assumed, due to high mitochondria copy number in the muscle tissue used for DNA extraction, that mitochondrial DNA was enriched in our WGS data. We used a minimum subset of 1.6 Gb raw PE data as input for *de novo* assembly per individual and a maximum of 4.8 Gb. Mitochondrial *de novo* assembly was carried out using the MitoZ toolkit v2.4 [81]. We used the ‘all’ function to assemble, annotate and plot genomes with parameters ‘genetic_code=5’ and ‘clade=Arthropoda’. In cases where assemblies yielded the reverse compliment of the mitogenome, identified by the gene order and orientation being the opposite to that of the ancestral insect [82], the mitogenome was derived by running the assembly through the reverse compliment function of MEGA-X v10.2.6 [83]. An additional 57 mitochondrial genomes were obtained from previous studies [20,25,26].

### Phylogenetic and population structure

A Maximum Likelihood (ML) phylogenetic tree was constructed using whole nuclear genomes to assess phylogenetic structure and relationships within the Black Soldier Fly species. We used RAxML program v8.2.9 [84] to construct a ML tree using the generalized time-reversible substitution model and gamma (GTRGAMMA) model, Lewis ascertainment bias and 100 bootstrap (BS) replications based on a random subset of 648,059 nuclear whole-genome SNPs. Further phylogenetic analysis using ASTRAL v5.7.8 [85], a coalescent based method robust to Incomplete Lineage Sorting (ILS), was performed to validate the ML analysis. A 10% sample of SNPs from each chromosome was extracted and split into 5,000 SNP segments and a gene tree was constructed on each using 100 BS replications totalling 8,800 trees, this was repeated for both the Neighbour-Joining (NJ) and ML method.

Phylogenetic reconstruction of the mitochondrial genomes was also performed. Whole mitochondrial genomes were aligned using MUSCLE v3.8.31 [86] and a phylogenetic ML tree produced using RAxML v8.2.9 [83]. A principal component analysis (PCA) [87] was performed using plink v1.9 [88] with Black Soldier Fly samples only. A random subsample of two individuals per unique geographic sampled population was used to conduct the PCA analysis after it was found that an unbalanced design artificially inflated genetic variation across both PC1 and PC2. Structure analysis was performed using ADMIXTURE v1.3.0 [89] assuming a minimum and maximum number of ancestral populations (*k*) of two and 11, respectively, based on the number of populations sampled. Linkage pruning was performed on SNPs used for both PCA and STRUCTURE analysis using plink v1.9 (--double-id --allow-extra-chr --set-missing-var-ids @:# --indep-pairwise 50 10 0.1) [88]. This filtering reduced the impact of non-independent SNPs reducing bias when estimating population structure ensuring sites were not disproportionately driven by physically linked loci. A NJ tree [90] was constructed using wild and captive Cytochrome Oxidase I (COI) mitochondrial marker genes from published and *de novo* assembled mitochondrial genomes. A NJ analysis of COI sequences was performed by aligning sequences with MUSCLE v3.8.31 [86] and phylogenetic analysis with MEGA v10.1.7 [83] using default parameters.

### Intraspecific divergence, diversity and genetic differentiation

Patterns of intraspecific divergence within the Black Soldier Fly complex were revealed using genome wide (nuclear) scans for genetic differentiation (*F_ST_*) and sequence divergence (*D_XY_*) on all populations. Non-overlapping 20 kb windows in 20 kb steps containing a minimum of 100 genotyped sites (-w 20000 -m 100 -s 20000) were used to generate genome-wide Nucleotide diversity (π), *F_ST_*and *D_XY_* statistics using popgenWindows.py (https://github.com/simonhmartin/genomics_general). Genome-wide nucleotide diversity was plotted in 20 kb windows using data smoothing (method = “loess”, span = 0.01).

### Divergence time analysis

Due to the paucity of Black Soldier Fly divergence data, a dated Diptera tree was first required to provide temporal calibration points for a Black Soldier Fly divergence analysis, as in Nardi *et al.* (2010) [91]. Here, we used mitochondrial DNA (mtDNA) only to estimate approximate divergences times. For the first calibration dating run mitogenomes of two Black Soldier Fly individuals, from divergent branches of both the nuclear and mitochondrial phylogeny and a single outgroup individual, *Ptecticus aurifer,* were obtained along with ten Diptera mitogenomes from GenBank (supplementary table 11) [92]. These dipteran mitochondrial genomes were aligned in MUSCLE v3.8.31 [86]. A molecular clock test was conducted on the resultant alignment and the optimal model of sequence evolution was selected by AIC using MEGA-X v10.2.6 [83]. A Bayesian dating analysis was conducted in BEAST v2.6.6 [93], using a GTR+G+I model of sequence evolution, and a relaxed lognormal molecular clock. Dating priors were set to calibrate six nodes spanning the Diptera phylogeny, based on fossil and biogeographic evidence, and the Yule model was used for the tree prior (supplementary table 4). Markov chain Monte Carlo (MCMC) length was set to 100 million, with samples logged every 1,000 steps. The model output was inspected in Tracer v1.7.2 [94]. After checking for parameter convergence, the burn-in was set to 25%, and the effective sample size of relevant parameters was verified to exceed 150.

A second BEAST run was conducted using temporal constraints derived in the first run for the Stratiomyidae and *H.illucens* nodes. This allowed estimation of key divergences within the Black Soldier Fly phylogeny, given the lack of molecular dating studies, individuals which spanned the phylogeny were used (supplementary table 4), alongside a *P. aurifer* outgroup. Alignment and divergence analysis was repeated a second time using the parameters previously described. We used a tree prior with a coalescent model of constant population size. Temporal constraints were placed on two nodes, using the output of the initial BEAST run, corresponding to the *P. aurifer* - Black Soldier Fly split and the common ancestor of the Black Soldier Fly clade (supplementary table 4). The MCMC chain was as previously described. After inspecting the model output in Tracer v1.7.2 [94], a burn-in of 10% was set. The dated phylogeny was derived using TreeAnnotator package of BEAST v2.6.6 [93]. The mitochondrial sequences used for both BEAST runs and their GenBank accession numbers are given in supplementary table 11, while all temporal constraints, outputs, and literature corroboration for phylogenetic nodes spanning both dating runs are given in supplementary table 4.

To corroborate mtDNA divergence time results we also estimated divergence times of Black Soldier Fly lineages using whole genome data. A random subsample of 3,978 genome-wide SNPs were used from all sequenced individuals for Bayesian dating analysis, again conducted in BEAST v2.7.7 [93]. Sex chromosome S7 was excluded from this analysis. A single temporal constraint was set for the *P. aurifer* and *H. illucens* split as our previous evidence and published evidence [95] were in agreement. A coalescent constant population model was used with the *Drosophila melanogaster* mutation rate of 2.8 x 10^-9^ as an estimate for the Black Soldier Fly mutation rate [96]. A starting tree was provided based on the phylogenetic tree produced in Figure 1A. A chain length of 10 million was used with a burn-in rate of 10% and sampling was performed every 1,000 chains. Our effective sample size was deemed suitable as it exceeded 4,000. Results were assessed and visualised in Tracer v1.7.2 [94].

### Genetic signals of domestication

Genomic diversity, and Linkage Disequilibrium (LD) decay were calculated to reveal genetic signatures of domestication in sampled populations with a sequencing depth >19x. Nucleotide diversity (π) and *F_ST_* was calculated in non-overlapping 100 kb windows using popgenWindows.py (-w 100000 -m 100 -s 100000) (https://github.com/simonhmartin/genomics_general) using 100 kb steps containing at least 100 genotyped sites. Values of π for every individual was presented as mean heterozygosity within each sampled population. Patterns of LD decay were first calculated genome wide for a random subset of three individuals for each population. We used PopLDdecay v3.41 [97] to generate pairwise LD *r*^2^ values setting a maximum distance of 50 kb between sampled SNPs.

To assess the impact of domestication in the Black Soldier Fly we performed scans for Runs of Homozygosity (RoH) to reveal genomic regions associated with positive selection and genetic drift. We tested each maintained captive population independently despite known shared population history and strong *k*-means clustering support for a single population structure of the Sheppard-lineage, due to a documented split approximately 1-2 decades ago. We also used patterns observed from *k*-means ancestry assignment (admixture) analysis to support our grouping into geographic clusters. All geographic populations were treated independently based off results at *k* = 11. Industry strains (A-C) were grouped based on population structure results and identical rearing conditions. Runs of Homozygosity were identified as in Generalovic *et al*. (2021) using the MAF (0.1) filtered VCF file with the following exceptions “windowSize = 20”, “minSNP = 20” and “minLengthBps = 250000” using the detectRUNS v0.98 R package [98]. Chromosome seven (X chromosome) and individuals with no detected homozygous regions were excluded from RoH analysis. Additional support using *F_ST_* with the closest sampled wild progenitor was acquired performed as in “Genetic signals of domestication”. Candidate loci were identified from gene codes in the GCA_905115235.1 reference by performing blastp searches against the Arthropoda blast nr database using an *e*-value threshold of 1*e*-5. Gene Ontology summary information for each candidate gene was obtained from flybase [99].

### Introgression and gene flow

Mito-nuclear discordance was predicted by plotting a co-phylogenetic plot of the previously constructed nuclear and mitochondrial phylogenies using the phytools v1.0-3 package [100] in R v3.6.3 [101]. Patterns of admixture were detected by revealing excess allele sharing between individuals using ancestral (ABBA) and derived (BABA) allele patterns [102,103], *f_dM_* values [104] and the *f-*branch statistic [105]. We used ABBA/BABA patterns using Dsuite v0.4r38 Dtrios function [106] and provided the previously resolved SNP based phylogeny (Figure 1A) as a tree file (-t). Trios showing *p*-values > 0.05 and Z-scores < 3 post *p*-value adjustment using Benjamini-Hochberg multiple correction in R v4.0.3 [102] were filtered out of the analysis. We generated the *f*-branch (*f_b_*) statistic using individuals sequenced to the highest coverage to represent populations within the phylogeny to assign gene flow to all, including internal ancestral, branches using *P. aurifer* as the outgroup [105]. Populations that were not monophyletic were represented by multiple individuals in the *f_b_* analysis. Genome-wide scans were performed on samples showing an excess of gene flow identified from the *f*-branch method using the Dsuite Dinvestigate function calculating average *f_dM_* over 50 SNPs in 25 SNP steps across the whole genome. Genomic hotspots of introgression were identified by setting an arbitrary 1% cut-off for *f_dM_*values.

## Acknowledgments

This work was supported by the Biotechnology and Biological Sciences Research Council (BB/M011194/1).

## Availability of supporting data

The data sets supporting the results of this article are available under Bioprojects PRJEB37575 and PRJEB58720.

## Notes

### Competing Interest Statement

The authors have declared no competing interest.

### Summary of Updates

This revision includes an update to the analysis removing selective sweep analysis and including analysis for runs of homozygosity. General text updates throughout included to tone down claims of global sampling. Figures 3 & 4 added in place of figure 3 (old).

