## supplementary figures for "Cryptic diversity and impacts of domestication in the Black Soldier Fly (*Hermetia illucens*) genome"

**Supplementary Figures:** **Cryptic diversity and signatures of domestication in the Black Soldier Fly (*Hermetia illucens*)**

Tomas N. Generalovic^1^*, Christoph Sandrock^2^, Benjamin J. Roberts^3^, Joana I. Meier^1,4^, Martin Hauser^5^, Ian A. Warren^1^, Miha Pipan^6^, Richard Durbin^7^ & Chris D. Jiggins^1^.

^1^Department of Zoology, University of Cambridge, Cambridge, UK; ^2^Department of Livestock Sciences, Research Institute of Organic Agriculture (FiBL), Frick, Switzerland; ^3^Georgina Mace Centre for the Living Planet, Faculty of Natural Sciences, Imperial College London, London, UK; ^4^Tree of Life Programme, Wellcome Sanger Institute, Wellcome Trust Genome Campus, Hinxton, Cambridge, UK; ^5^California Department of Food and Agriculture, Plant Pest Diagnostics Branch, Sacramento, CA, USA; ^6^Better Origin, Entomics Biosystems Limited, Cambridge, UK; ^7^Department of Genetics, University of Cambridge, Cambridge, UK

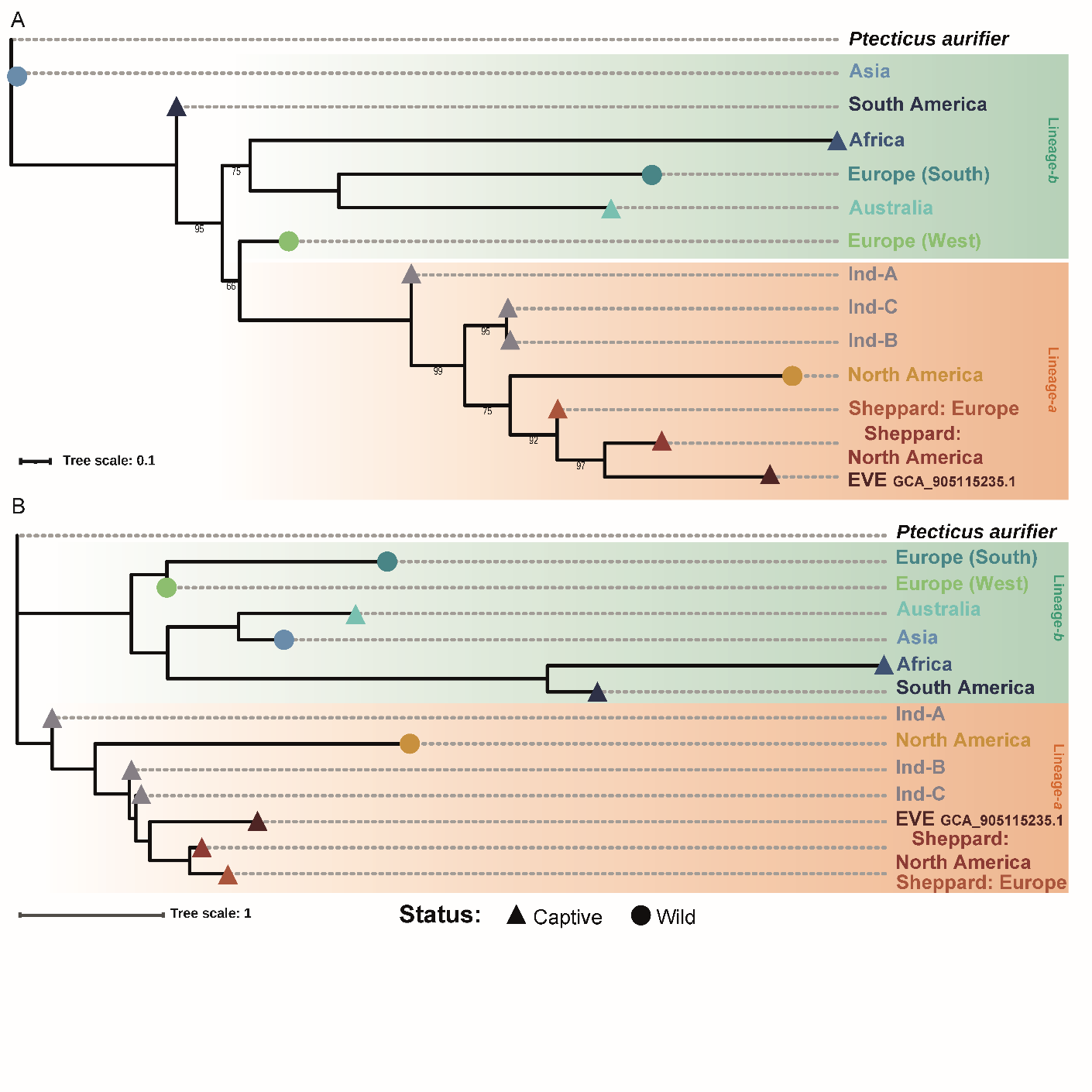

**Supplementary figure 1.** Phylogenetic analysis of the Black Soldier Fly using the ASTRAL coalescent method for both Neighbor-Joining (NJ) (A) and Maximum-Likelihood (ML) (B) analysis. Species phylogenies are generated using 88 gene trees of regions containing 5,000 SNPs and 100 bootstrap replications each, totaling 8,800 gene trees. NJ analysis indicates the Asian population is closest to the outgroup, a species found in Asia, whereas South America remains basal to the remaining Black Soldier Fly lineages. Additionally, the western European population now appears as the source to lineage-*a* (A). ML species tree using ASTRAL confirms the same pattern observed with the larger dataset presented in the manuscript (B). Bootstrap values lower than 100 are annotated, fully supported branches are unlabeled.

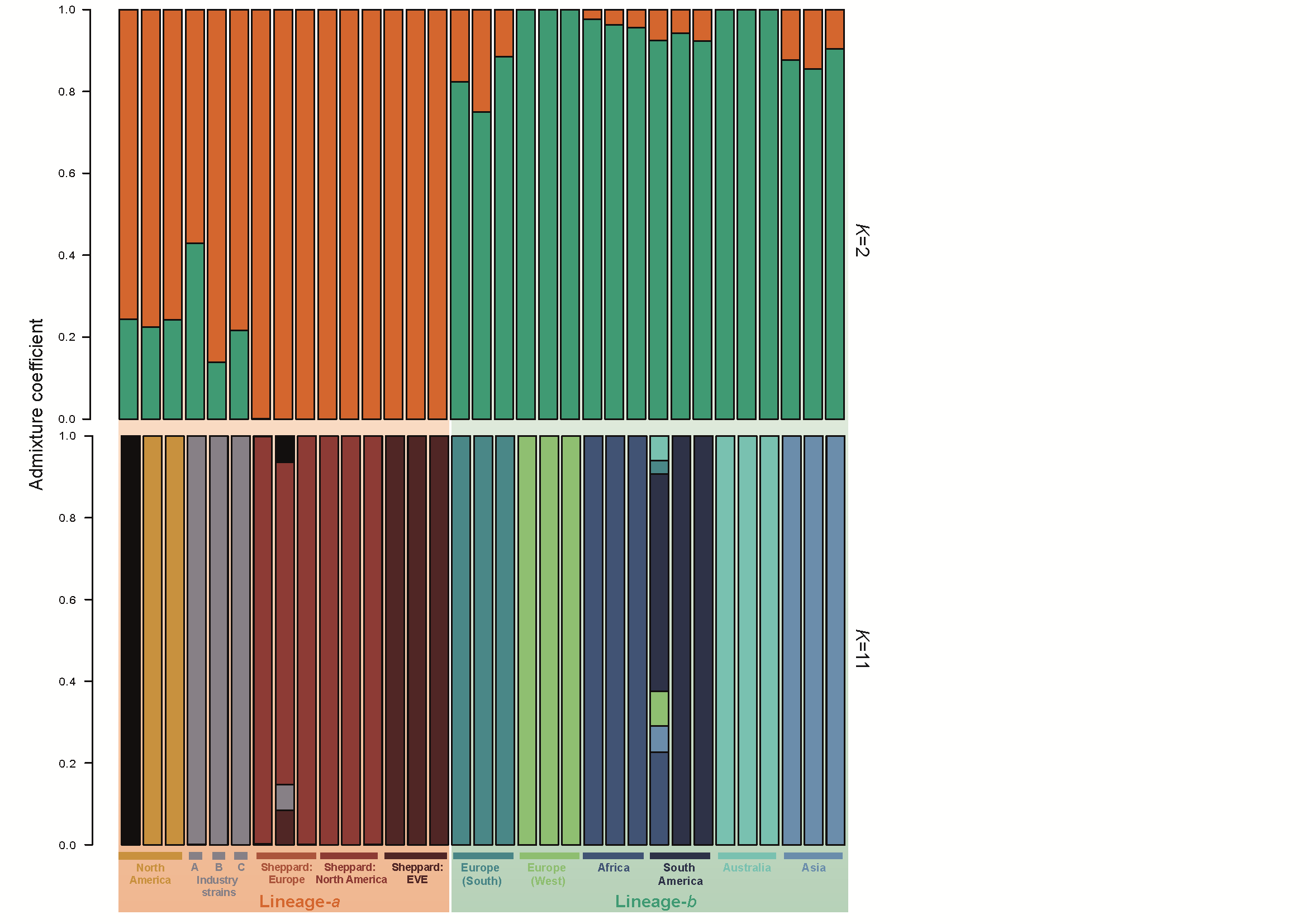

**Supplementary figure 2.** ADMIXTURE analysis of Black Soldier Fly individuals from sampled populations. Sample dataset sub-sampled down to three of the highest sequenced individuals per population. Repeated analysis of ancestry *k*-means clustering using a reduced and balanced sample size recapitulates key results when utilizing all individuals (Figure 1B). However, we observe that the Sheppard strain becomes collapsed into one ancestry as predicted due to a recent documented split of these two populations. Conversely, we observe individuals from wild North America being assigned to two distinct clusters.

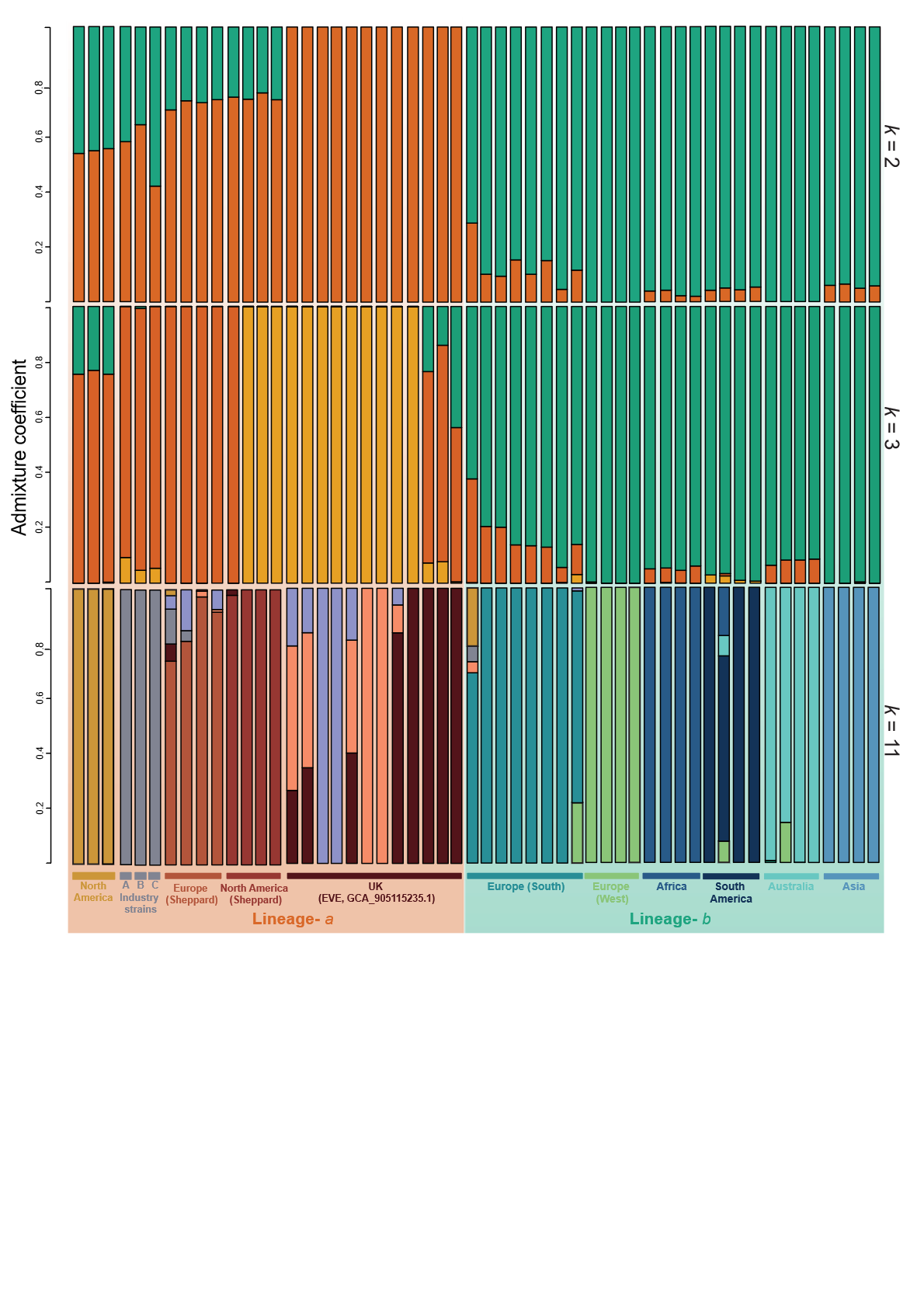

**Supplementary figure 3.** ADMIXTURE analysis of Black Soldier Fly individuals from sampled populations. Analysis of ancestry *k*-means clustering including *k* = 2 (0.885), *k* = 3 (0.887) and *k* = 11 (1.764). A two-lineage hypothesis is supported at *k* = 2, however, at *k* = 3 we observe that lineage-*a* captive populations EVE and North America predominately make up this third cluster. The remaining populations are clustered within previously identified lineage-*a* and -*b* at this *k* = 3 analysis. Results of *k* = 11 provided for context as in Figure 1.

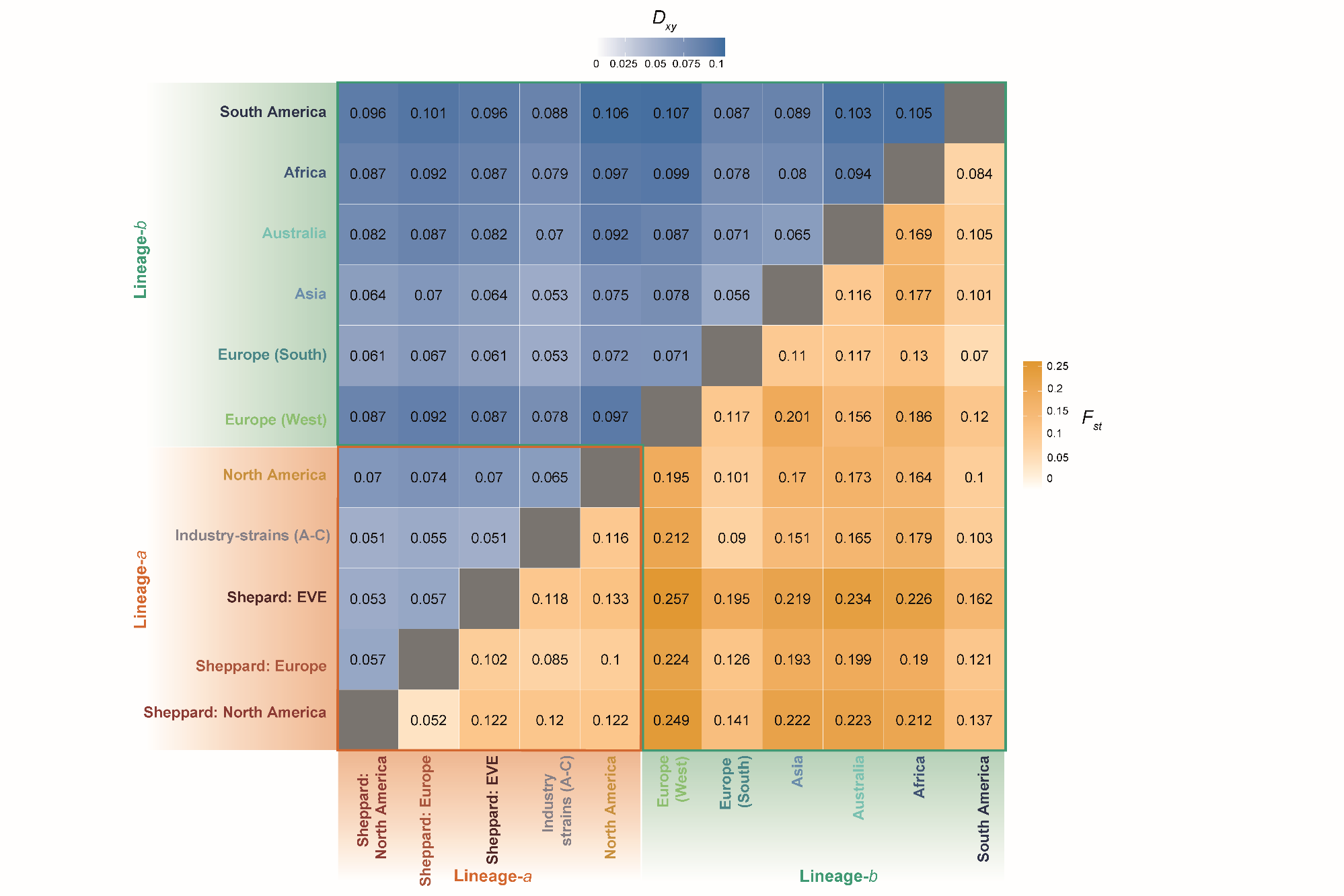

**Supplementary figure 4.** Pairwise genetic differentiation (*F_ST_*) and absolute sequence divergence (*d_XY_*) matrix of Black Soldier Fly individuals from sampled global populations belonging to the two major clades. All 54 individuals were grouped within their sampling populations as indicated in Figure 1A&B, and all three industry strains A-C were treated as a single population according to respective assignments of the admixture analysis.

**Supplementary figure 5.** Mitochondrial genome assembly of the Black Soldier Fly "EVE" GCA_905115235.1 reference-line using individual CAM006158 as a plotted example. Assembled mitochondrial genome circos plot is annotated to show gene order (A), sequence coverage (B) and GC content (C).

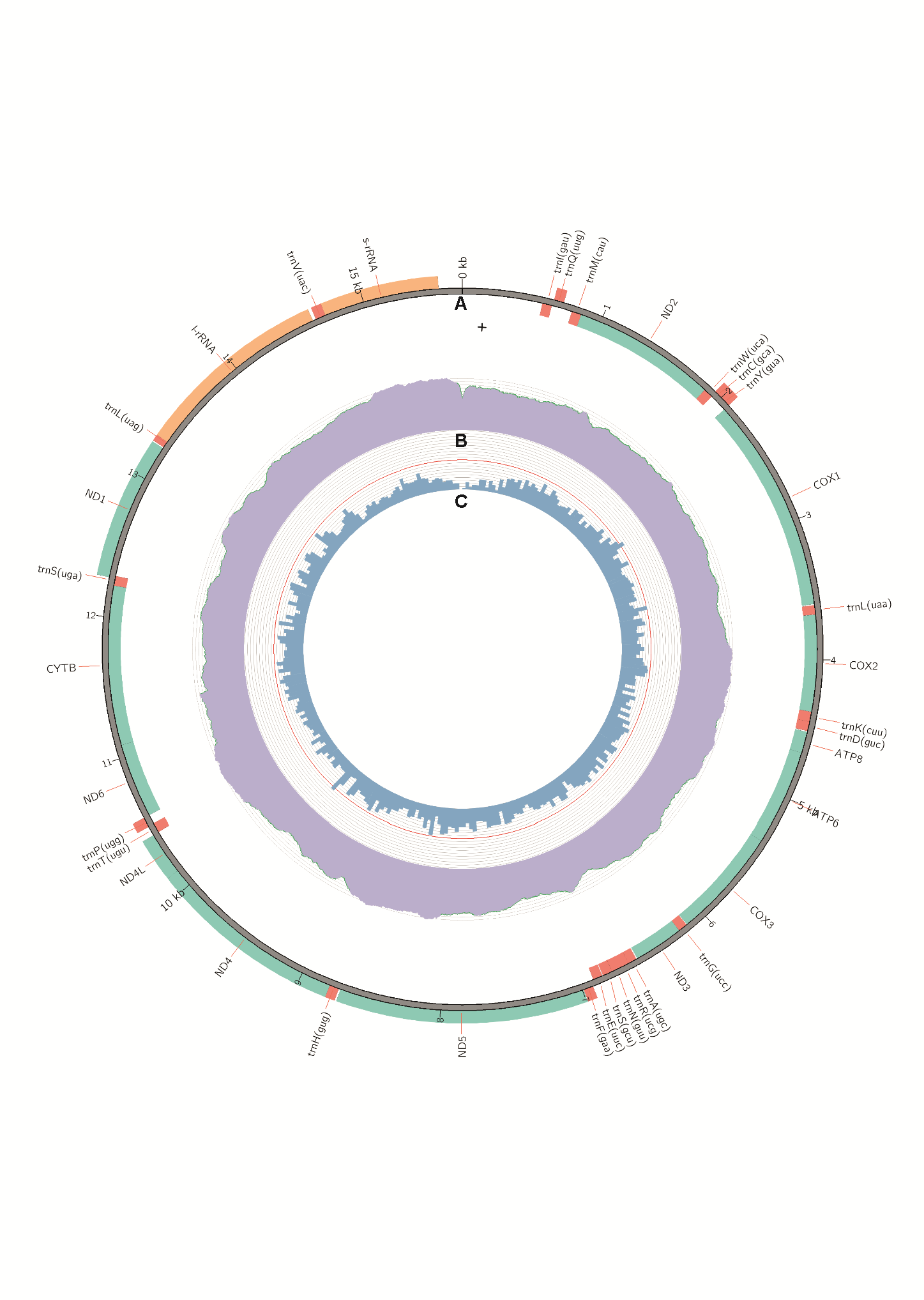

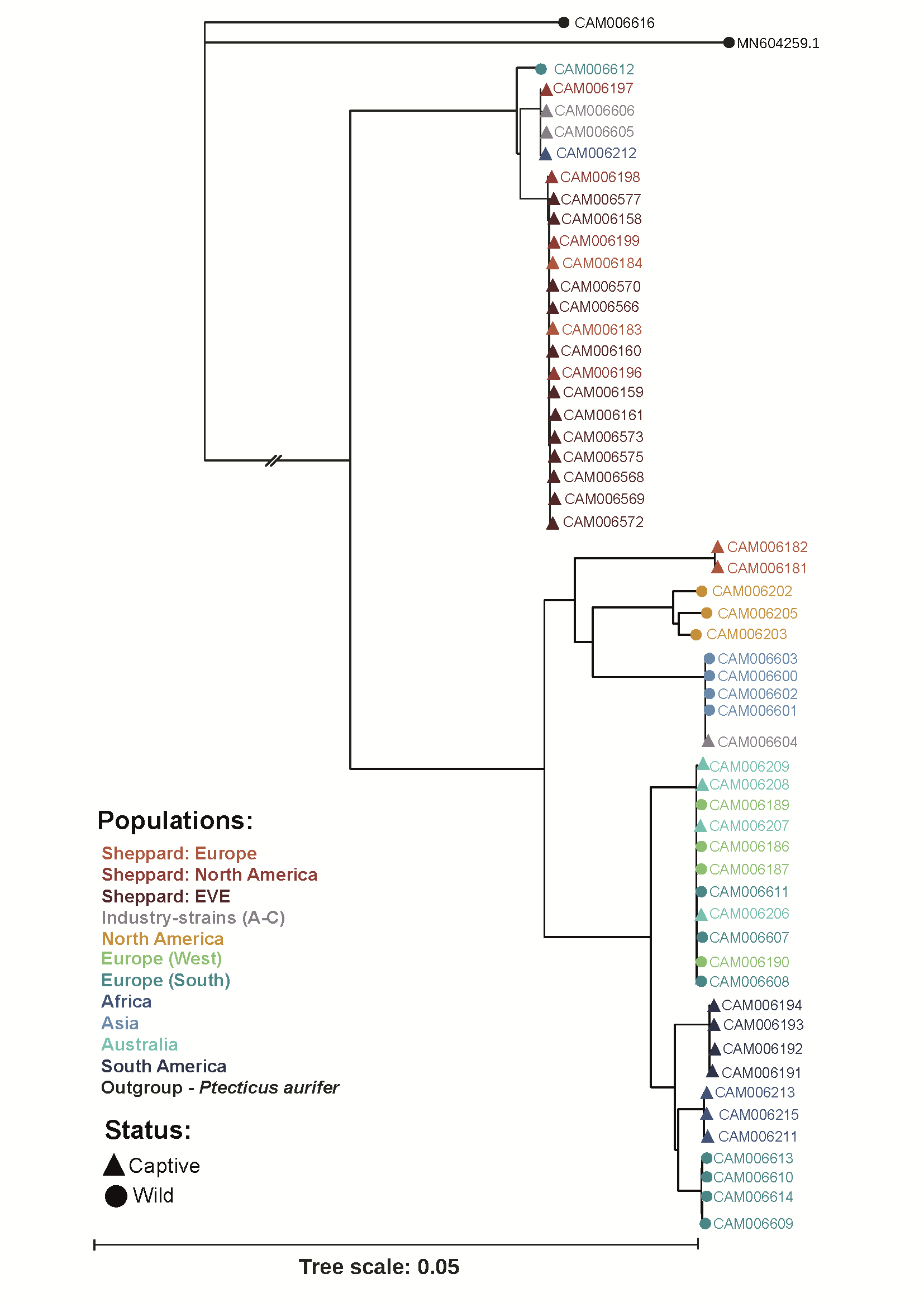

**Supplementary figure 6.** Maximum Likelihood phylogenetic reconstruction using whole mitochondrial genomes of Black Soldier Fly (n=54) and *Ptecticus aurifer* (n=2). Outgroup species *P. aurifer* produced *de novo* in this study (CAM006616) clusters with previously published MN604259.1 individual (Zhan *et al*., 2020).

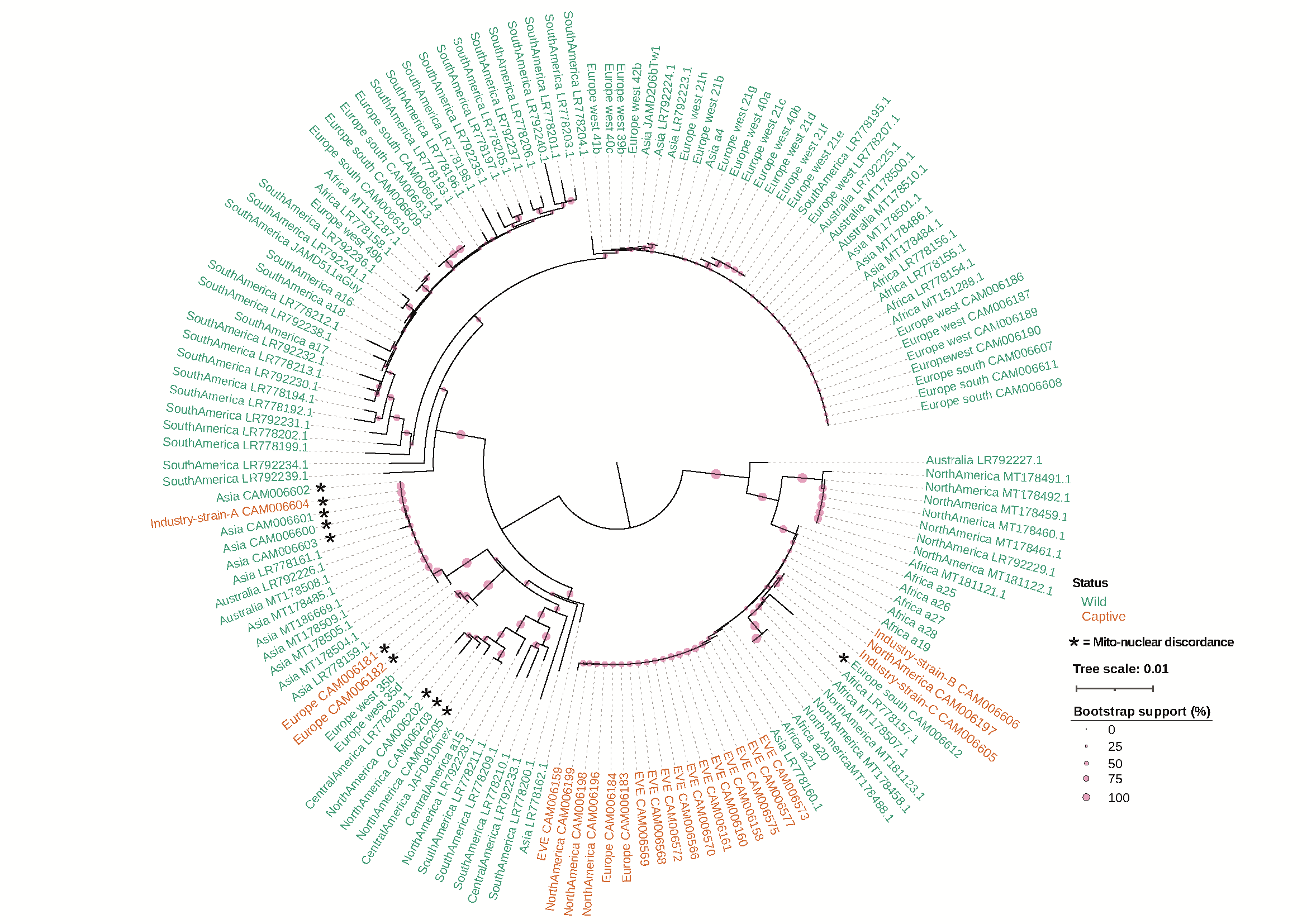

**Supplementary figure 7.** Neighbor-Joining (NJ) phylogeny using a sub-sample of Black Soldier Fly cytochrome oxidase I (COI) markers produced in this (CAM identifiers) and previously published (Guiliet *et al*., 2021; Stahls *et al*., 2019) studies. A sample of globally dispersed populations are represented within this tree and comprised of several wild (green) and extensively reared captive (captive; orange) samples, as labelled by their continental or strain origins. Samples presenting patterns of mito-nuclear discordance from previous whole nuclear genome analysis (exclusively applicable to CAM identifiers) are indicated using an asterisk. Bootstrap support for the NJ phylogeny is annotated on tree nodes. Phylogenetic NJ tree shows strong support for a wild North American origin for the captive populations of the Black Soldier Fly with evidence of mito-nuclear discordance explaining sample locations outside of this lineage-*a* clade.

**Supplementary figure 8.** Whole-genome divergence analysis using a subset of genome wide SNP alignments (*n* = 3,978) for the phylogenetic tree previously identified on whole genome analysis. Compared to mtDNA divergence analysis the previously published a1_Ghana sample from Gulliet et al (2021) is excluded due to lacking nuclear data.

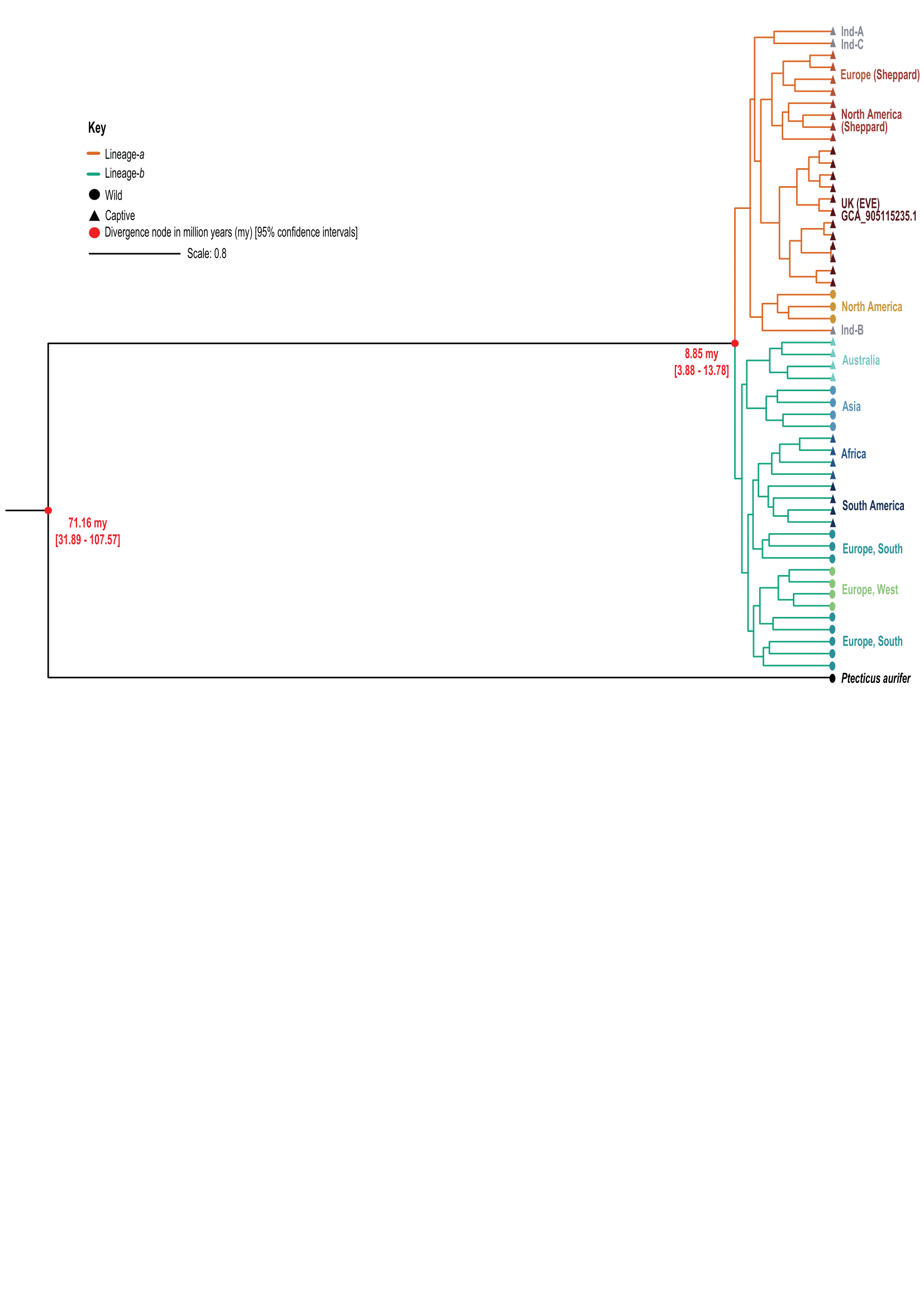

**Supplementary figure 9.** Genome-wide heterozygosity levels of Black Soldier Fly populations above 19x sequence coverage.

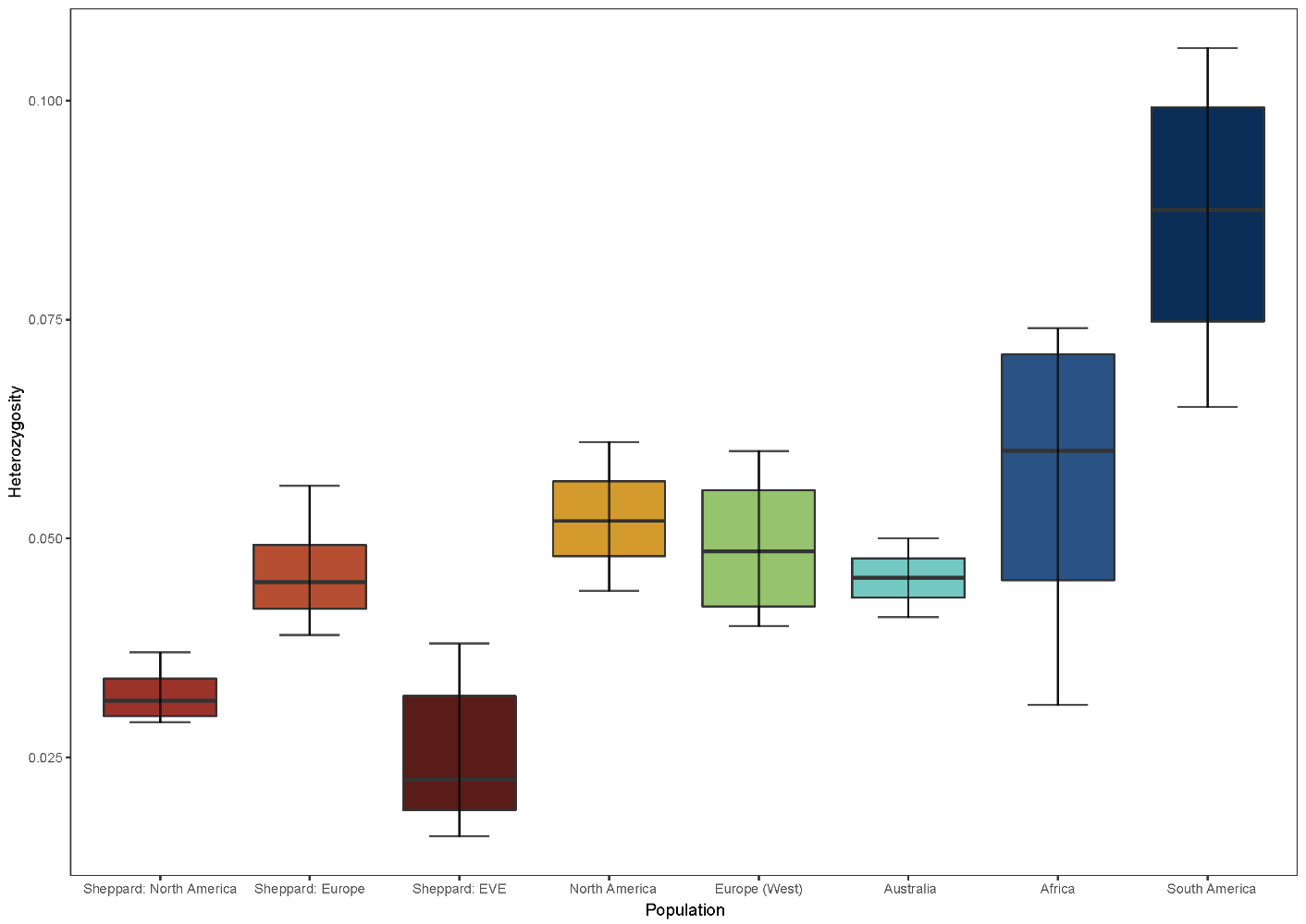

**Supplementary figure 10.** Estimates of linkage-disequilibrium breakdown of captive and wild Black Soldier Fly populations measured over 50 kb. Populations with adequate sequencing depth as in supplementary figure 9 are included.

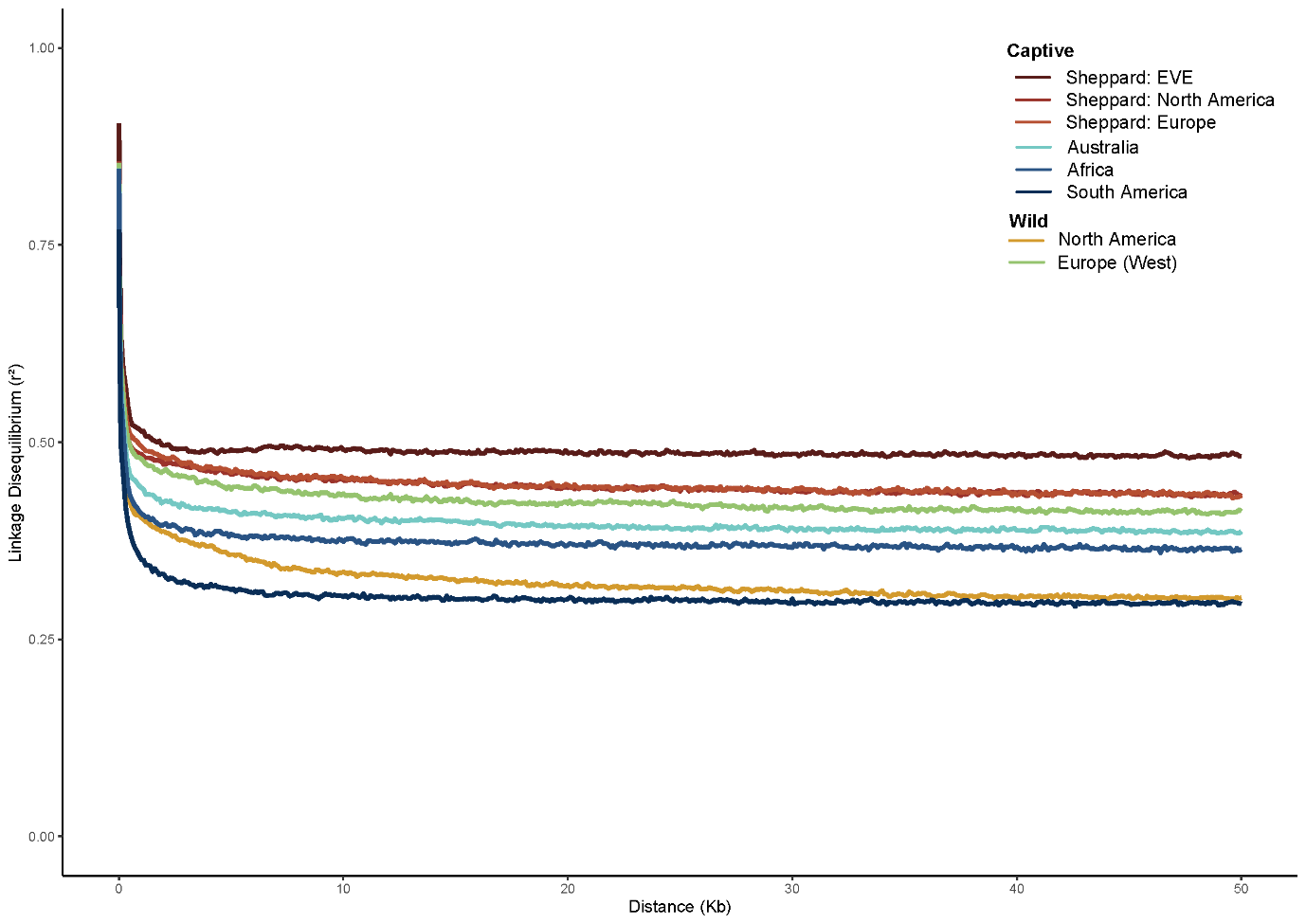

**Supplementary figure 11.** Shared, captive and wild only Runs of Homozygosity (RoH) within the sampled Black Soldier Fly genomes. Total number of RoH, mean length of RoH and size category for long or short RoH indicated for each captivity status assessed including shared regions.

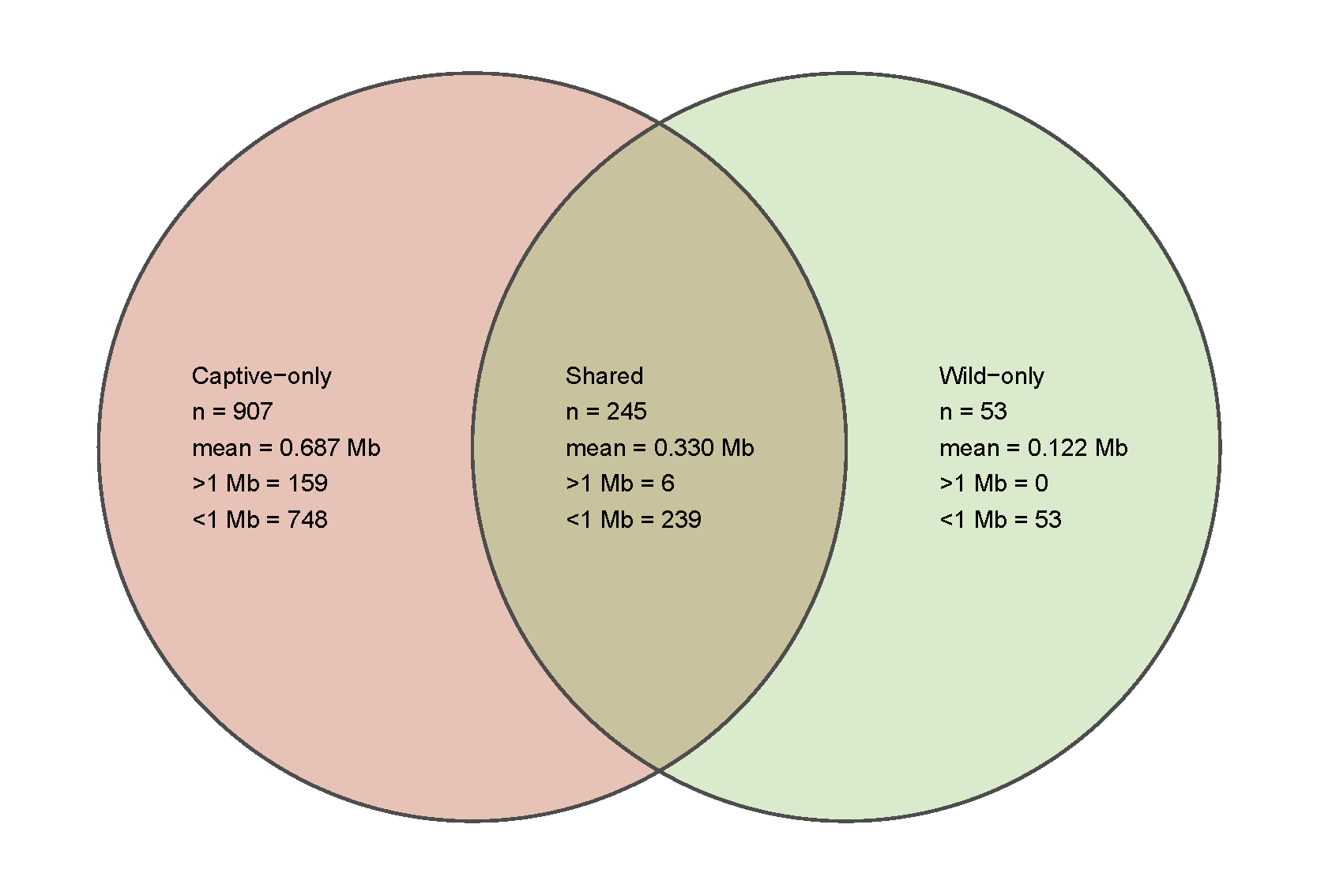

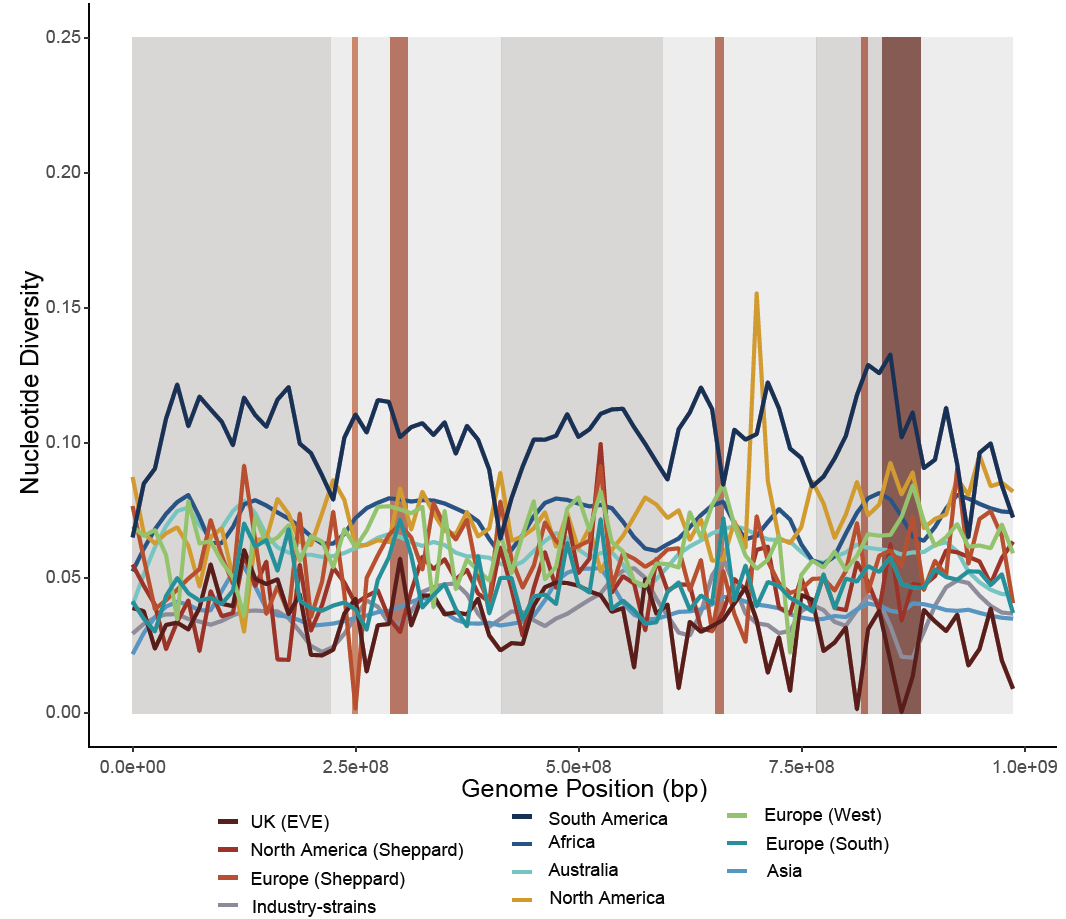

**Supplementary figure 12.** Genome-wide nucleotide diversity (*π*) averaged for all sampled populations. Population specific colour is provided along for “smoothed” data lines due to high noise levels of raw data. Chromosomes are indicated in background alternating blocks and the five major regions associated with domestication are provided across the identified sweep candidate co-ordinates.

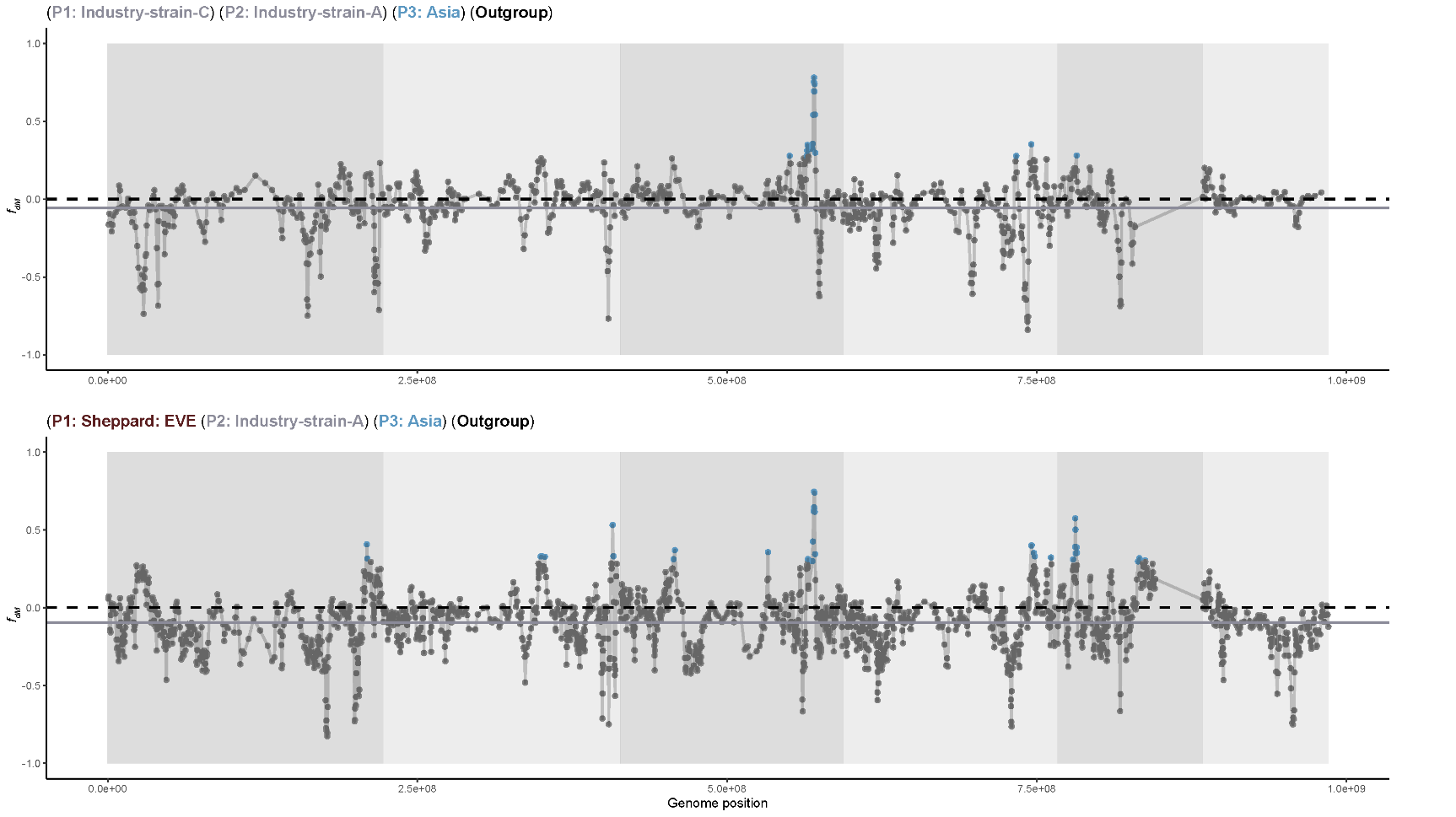

**Supplementary figure 13.** Genome-wide *f_dM_* of the Black Soldier Fly highlighting 99^th^ percentile outlier values of allele sharing between the wild Asian population (P3) representative and industry-strain-A (Ind-A; P2). Mean genome-wide levels of *f_dM_* in Ind-A are indicated by a solid grey line. Assumptions of no gene flow, the null hypothesis, is presented as a black dashed line. Strong signals (top 1%/ 99^th^ percentile) of wild Asian introgression into P2 are presented as blue points.
