## supplementary tables for "Cryptic diversity and impacts of domestication in the Black Soldier Fly (*Hermetia illucens*) genome"

**Supplementary Tables:** **Cryptic diversity and signatures of domestication in the Black Soldier Fly (*Hermetia illucens*)**

Tomas N. Generalovic^1^*, Christoph Sandrock^2^, Benjamin J. Roberts^3^, Joana I. Meier^1,4^, Martin Hauser^5^, Ian A. Warren^1^, Miha Pipan^6^, Richard Durbin^7^ & Chris D. Jiggins^1^.

^1^Department of Zoology, University of Cambridge, Cambridge, UK; ^2^Department of Livestock Sciences, Research Institute of Organic Agriculture (FiBL), Frick, Switzerland; ^3^Georgina Mace Centre for the Living Planet, Faculty of Natural Sciences, Imperial College London, London, UK; ^4^Tree of Life Programme, Wellcome Sanger Institute, Wellcome Trust Genome Campus, Hinxton, Cambridge, UK; ^5^California Department of Food and Agriculture, Plant Pest Diagnostics Branch, Sacramento, CA, USA; ^6^Better Origin, Entomics Biosystems Limited, Cambridge, UK; ^7^Department of Genetics, University of Cambridge, Cambridge, UK

| **Supplementary table 1.** Sample information and sequencing statistics. Sample information is provided for each sequenced whole genome with geographic and sample source. Genome mapping, to the *Hermetia illucens* reference genome as in Generalovic et al. (2021), coverage is predicted assuming a genome size of 1.01 Gb. All samples are *Hermetia illucens* species except for CAM006616 which is *Ptecticus aurifer*. | | | | | | | |
| --- | --- | --- | --- | --- | --- | --- | --- |
| **Sample ID (CAMID)** | **Geographic position** | **Captivity status** | **Source** | **Sex** | **Mapped reads (#)** | **Mapped reads (%)** | **Mapped coverage (x)** |
| CAM006196 | North America | Captive | Kaya *et al*., 2021 | Male | 161,800,218 | 96.25 | 23.1 |
| CAM006197 | North America | Captive | Kaya *et al*., 2021 | Female | 163,224,078 | 65.15 | 20.8 |
| CAM006198 | North America | Captive | Kaya *et al*., 2021 | Female | 166,050,414 | 96.24 | 23.1 |
| CAM006199 | North America | Captive | Kaya *et al*., 2021 | Female | 165,547,061 | 96.17 | 23.5 |
| CAM006181 | Europe | Captive | Kaya *et al*., 2021 | Female | 252,775,923 | 96.54 | 35.4 |
| CAM006182 | Europe | Captive | Kaya *et al*., 2021 | Male | 180,127,723 | 96.64 | 24.8 |
| CAM006183 | Europe | Captive | Kaya *et al*., 2021 | Female | 172,112,061 | 96.78 | 24.3 |
| CAM006184 | Europe | Captive | Kaya *et al*., 2021 | Male | 181,009,193 | 96.82 | 25.4 |
| CAM006158 | UK | Captive | Generalovic et al., 2021 | Female | 166,453,262 | 96.67 | 23.7 |
| CAM006159 | UK | Captive | Generalovic et al., 2021 | Male | 163,262,946 | 96.52 | 23.2 |
| CAM006160 | UK | Captive | Generalovic et al., 2021 | Female | 165,012,025 | 96.62 | 23.4 |
| CAM006161 | UK | Captive | Generalovic et al., 2021 | Male | 163,752,443 | 96.53 | 23.3 |
| CAM006570 | UK | Captive | Generalovic et al., 2021 | Female | 163,879,564 | 96.31 | 23.8 |
| CAM006566 | UK | Captive | Generalovic et al., 2021 | Female | 160,569,238 | 96.41 | 23.3 |
| CAM006572 | UK | Captive | Generalovic et al., 2021 | Female | 164,233,561 | 96.43 | 23.6 |
| CAM006568 | UK | Captive | Generalovic et al., 2021 | Female | 151,263,233 | 96.43 | 21.8 |
| CAM006569 | UK | Captive | Generalovic et al., 2021 | Male | 163,109,345 | 96.34 | 23.5 |
| CAM006573 | UK | Captive | Generalovic et al., 2021 | Female | 163,806,471 | 96.56 | 23.4 |
| CAM006577 | UK | Captive | Generalovic et al., 2021 | Male | 163,910,656 | 96.65 | 23.4 |
| CAM006575 | UK | Captive | Generalovic et al., 2021 | Male | 161,327,841 | 96.59 | 23.2 |
| CAM006604 | Industry-strain-A (Asia source, UK reared) | Captive | Better Origin | Male | 123,199,074 | 95.04 | 17.5 |
| CAM006605 | Industry-strain-C, (Asia source, UK reared) | Captive | Better Origin | Male | 112,136,879 | 95.31 | 15.8 |
| CAM006606 | Industry-strain-B, (Asia source, UK reared) | Captive | Better Origin | Male | 111,673,733 | 95.76 | 15.5 |
| CAM006202 | North America | Wild | Kaya *et al*., 2021 | Male | 165,907,749 | 67.25 | 22.6 |
| CAM006203 | North America | Wild | Kaya *et al*., 2021 | Female | 182,664,875 | 95.11 | 24.8 |
| CAM006205 | North America | Wild | Kaya *et al*., 2021 | Male | 188,010,026 | 96.16 | 26.2 |
| CAM006603 | Asia | Wild | Martin Hauser (CDFA) | Female | 118,385,405 | 96.61 | 16.5 |
| CAM006600 | Asia | Wild | Martin Hauser (CDFA) | Male | 85,950,287 | 96.69 | 12.2 |
| CAM006601 | Asia | Wild | Martin Hauser (CDFA) | Female | 136,816,978 | 95.47 | 19 |
| CAM006602 | Asia | Wild | Martin Hauser (CDFA) | Male | 82,078,666 | 96.80 | 11.8 |
| CAM006186 | Europe (west) | Wild | Kaya *et al*., 2021 | Female | 167,631,259 | 94.91 | 23.3 |
| CAM006187 | Europe (west) | Wild | Kaya *et al*., 2021 | Female | 187,581,181 | 95.44 | 26.3 |
| CAM006189 | Europe (west) | Wild | Kaya *et al*., 2021 | Male | 182,960,993 | 95.22 | 25.6 |
| CAM006190 | Europe (west) | Wild | Kaya *et al*., 2021 | Male | 164,509,454 | 94.26 | 23.3 |
| CAM006614 | Europe (south) | Wild | Better Origin | Male | 113,994,373 | 94.59 | 16 |
| CAM006613 | Europe (south) | Wild | Better Origin | Male | 114,380,634 | 94.45 | 16 |
| CAM006612 | Europe (south) | Wild | Better Origin | Male | 115,312,808 | 95.25 | 16 |
| CAM006611 | Europe (south) | Wild | Better Origin | Male | 138,780,503 | 94.66 | 19.5 |
| CAM006610 | Europe (south) | Wild | Better Origin | Male | 116,385,321 | 94.70 | 16.3 |
| CAM006609 | Europe (south) | Wild | Better Origin | Male | 114,108,446 | 94.72 | 15.9 |
| CAM006607 | Europe (south) | Wild | Better Origin | Male | 114,872,429 | 94.71 | 16.1 |
| CAM006608 | Europe (south) | Wild | Better Origin | Male | 116,946,151 | 94.71 | 16.4 |
| CAM006211 | Africa | Captive | Kaya *et al*., 2021 | Male | 191,008,044 | 94.42 | 26.6 |
| CAM006212 | Africa | Captive | Kaya *et al*., 2021 | Male | 134,908,203 | 79.17 | 19 |
| CAM006213 | Africa | Captive | Kaya *et al*., 2021 | Female | 159,984,901 | 98.09 | 22.8 |
| CAM006215 | Africa | Captive | Kaya *et al*., 2021 | Male | 184,505,155 | 95.97 | 25.8 |
| CAM006191 | South America | Captive | Kaya *et al*., 2021 | Female | 234,148,745 | 94.98 | 32 |
| CAM006192 | South America | Captive | Kaya *et al*., 2021 | Male | 161,854,374 | 95.02 | 22.8 |
| CAM006193 | South America | Captive | Kaya *et al*., 2021 | Male | 233,379,595 | 94.94 | 32.1 |
| CAM006194 | South America | Captive | Kaya *et al*., 2021 | Female | 182,142,300 | 95.49 | 25.1 |
| CAM006206 | Australia | Captive | Kaya *et al*., 2021 | Female | 160,625,708 | 94.44 | 22.5 |
| CAM006207 | Australia | Captive | Kaya *et al*., 2021 | Male | 161,200,325 | 94.49 | 22.6 |
| CAM006208 | Australia | Captive | Kaya *et al*., 2021 | Female | 179,607,215 | 95.80 | 25.2 |
| CAM006209 | Australia | Captive | Kaya *et al*., 2021 | Female | 170,352,448 | 94.50 | 24 |
| CAM006616 | Asia | Wild | Martin Hauser (CDFA) | NA | 45,791,226 | 51.24 | NA |

| **Supplementary table 2.** Cross Validation (CV) error rate output from ADMIXTURE analysis. | |
| --- | --- |
| ***k*-value** | **Cross Validation error (CV)** |
| 2 | 0.885 |
| 3 | 0.887 |
| 4 | 0.961 |
| 5 | 1.006 |
| 6 | 1.101 |
| 7 | 1.232 |
| 8 | 1.350 |
| 9 | 1.520 |
| 10 | 1.551 |
| 11 | 1.764 |

| **Supplementary table 3.** Mitochondrial (mt) assembly and annotation statistics. Sample information is also provided for each *de novo* whole mitochondrial assembly. Whole genome sequencing (WGS) was performed for each individual from muscle tissue allowing mtDNA enrichment in WGS data. We performed mt assembly using a subset of the WGS data. All samples are *Hermetia illucens* species except for CAM006616 which is *Ptecticus aurifer*. | | | | | | | | | | |
| --- | --- | --- | --- | --- | --- | --- | --- | --- | --- | --- |
| **Sample ID (CAMID)** | **Geographic Position** | **Captivity Status** | **Size (bp; primary contig)** | **%GC** | **%AT** | **GC-Skew** | **AT-Skew** | **Protein coding genes** | **tRNA genes** | **rRNA genes** |
| CAM006196 | North America | Captive | 15,698 | 0.281 | 0.719 | -0.251 | 0.014 | 13 | 22 | 2 |
| CAM006197 | North America | Captive | 15,696 | 0.282 | 0.718 | -0.254 | 0.014 | 13 | 22 | 2 |
| CAM006198 | North America | Captive | 15,698 | 0.281 | 0.719 | -0.251 | 0.014 | 13 | 22 | 2 |
| CAM006199 | North America | Captive | 15,696 | 0.281 | 0.719 | -0.251 | 0.014 | 13 | 22 | 2 |
| CAM006181 | Europe | Captive | 15,689 | 0.280 | 0.720 | -0.262 | 0.018 | 13 | 22 | 2 |
| CAM006182 | Europe | Captive | 15,688 | 0.280 | 0.720 | -0.262 | 0.018 | 13 | 22 | 2 |
| CAM006183 | Europe | Captive | 15,697 | 0.281 | 0.719 | -0.251 | 0.014 | 13 | 22 | 2 |
| CAM006184 | Europe | Captive | 15,696 | 0.281 | 0.719 | -0.251 | 0.014 | 13 | 22 | 2 |
| CAM006158 | UK | Captive | 15,696 | 0.281 | 0.719 | -0.251 | 0.014 | 13 | 22 | 2 |
| CAM006159 | UK | Captive | 15,822 | 0.280 | 0.720 | -0.252 | 0.012 | 13 | 22 | 2 |
| CAM006160 | UK | Captive | 15,697 | 0.281 | 0.719 | -0.251 | 0.014 | 13 | 22 | 2 |
| CAM006161 | UK | Captive | 15,822 | 0.280 | 0.720 | -0.252 | 0.012 | 13 | 22 | 2 |
| CAM006570 | UK | Captive | 15,696 | 0.281 | 0.719 | -0.251 | 0.014 | 13 | 22 | 2 |
| CAM006566 | UK | Captive | 15,696 | 0.281 | 0.719 | -0.251 | 0.014 | 13 | 22 | 2 |
| CAM006572 | UK | Captive | 15,698 | 0.281 | 0.719 | -0.251 | 0.014 | 13 | 22 | 2 |
| CAM006568 | UK | Captive | 15,697 | 0.281 | 0.719 | -0.251 | 0.014 | 13 | 22 | 2 |
| CAM006569 | UK | Captive | 15,697 | 0.281 | 0.719 | -0.251 | 0.014 | 13 | 22 | 2 |
| CAM006573 | UK | Captive | 15,697 | 0.281 | 0.719 | -0.251 | 0.014 | 13 | 22 | 2 |
| CAM006577 | UK | Captive | 15,696 | 0.281 | 0.719 | -0.251 | 0.014 | 13 | 22 | 2 |
| CAM006575 | UK | Captive | 15,697 | 0.281 | 0.719 | -0.251 | 0.014 | 13 | 22 | 2 |
| CAM006604 | Industry-strain, UK | Captive | 15,675 | 0.282 | 0.718 | -0.251 | 0.017 | 13 | 22 | 2 |
| CAM006605 | Industry-strain, UK | Captive | 15,694 | 0.282 | 0.718 | -0.255 | 0.013 | 13 | 22 | 2 |
| CAM006606 | Industry-strain, UK | Captive | 15,695 | 0.282 | 0.718 | -0.254 | 0.015 | 13 | 22 | 2 |
| CAM006202 | North America | Wild | 15,672 | 0.281 | 0.719 | -0.256 | 0.018 | 13 | 22 | 2 |
| CAM006203 | North America | Wild | 15,673 | 0.282 | 0.718 | -0.254 | 0.018 | 13 | 22 | 2 |
| CAM006205 | North America | Wild | 15,669 | 0.282 | 0.718 | -0.254 | 0.018 | 13 | 22 | 2 |
| CAM006603 | Asia | Wild | 13,862 | 0.294 | 0.706 | -0.240 | 0.020 | 13 | 21 | 1 |
| CAM006600 | Asia | Wild | 15,500 | 0.285 | 0.715 | -0.251 | 0.017 | 13 | 22 | 2 |
| CAM006601 | Asia | Wild | 15,537 | 0.285 | 0.715 | -0.251 | 0.017 | 13 | 22 | 2 |
| CAM006602 | Asia | Wild | 15,515 | 0.285 | 0.715 | -0.251 | 0.017 | 13 | 22 | 2 |
| CAM006186 | Europe (west) | Wild | 15,669 | 0.284 | 0.716 | -0.256 | 0.017 | 13 | 22 | 2 |
| CAM006187 | Europe (west) | Wild | 15,670 | 0.284 | 0.716 | -0.256 | 0.017 | 13 | 22 | 2 |
| CAM006189 | Europe (west) | Wild | 15,565 | 0.285 | 0.715 | -0.255 | 0.018 | 13 | 22 | 2 |
| CAM006190 | Europe (west) | Wild | 15,741 | 0.283 | 0.717 | -0.256 | 0.015 | 13 | 22 | 2 |
| CAM006614 | Europe (south) | Wild | 15,740 | 0.282 | 0.718 | -0.257 | 0.015 | 13 | 22 | 2 |
| CAM006613 | Europe (south) | Wild | 15,672 | 0.283 | 0.717 | -0.257 | 0.017 | 13 | 22 | 2 |
| CAM006612 | Europe (south) | Wild | 15,698 | 0.282 | 0.718 | -0.252 | 0.013 | 13 | 22 | 2 |
| CAM006611 | Europe (south) | Wild | 15,555 | 0.286 | 0.714 | -0.255 | 0.017 | 13 | 22 | 2 |
| CAM006610 | Europe (south) | Wild | 15,671 | 0.283 | 0.717 | -0.257 | 0.017 | 13 | 22 | 2 |
| CAM006609 | Europe (south) | Wild | 15,741 | 0.282 | 0.718 | -0.257 | 0.015 | 13 | 22 | 2 |
| CAM006607 | Europe (south) | Wild | 15,498 | 0.286 | 0.714 | -0.255 | 0.018 | 13 | 22 | 2 |
| CAM006608 | Europe (south) | Wild | 15,742 | 0.283 | 0.717 | -0.256 | 0.015 | 13 | 22 | 2 |
| CAM006211 | Africa | Captive | 15,745 | 0.282 | 0.718 | -0.256 | 0.014 | 13 | 22 | 2 |
| CAM006212 | Africa | Captive | 15,695 | 0.282 | 0.718 | -0.254 | 0.014 | 13 | 22 | 2 |
| CAM006213 | Africa | Captive | 15,671 | 0.283 | 0.717 | -0.257 | 0.017 | 13 | 22 | 2 |
| CAM006215 | Africa | Captive | 15,574 | 0.284 | 0.716 | -0.256 | 0.018 | 13 | 22 | 2 |
| CAM006191 | South America | Captive | 15,573 | 0.284 | 0.716 | -0.256 | 0.017 | 13 | 22 | 2 |
| CAM006192 | South America | Captive | 15,573 | 0.284 | 0.716 | -0.256 | 0.017 | 13 | 22 | 2 |
| CAM006193 | South America | Captive | 15,573 | 0.284 | 0.716 | -0.256 | 0.017 | 13 | 22 | 2 |
| CAM006194 | South America | Captive | 15,574 | 0.284 | 0.716 | -0.256 | 0.017 | 13 | 22 | 2 |
| CAM006206 | Australia | Captive | 15,741 | 0.284 | 0.716 | -0.257 | 0.016 | 13 | 22 | 2 |
| CAM006207 | Australia | Captive | 15,669 | 0.284 | 0.716 | -0.256 | 0.017 | 13 | 22 | 2 |
| CAM006208 | Australia | Captive | 15,271 | 0.287 | 0.713 | -0.252 | 0.018 | 13 | 22 | 2 |
| CAM006209 | Australia | Captive | 15,671 | 0.284 | 0.716 | -0.256 | 0.017 | 13 | 22 | 2 |
| CAM006616 | Asia | Wild | 15,346 | 0.267 | 0.733 | -0.203 | 0.000 | 13 | 22 | 2 |

| **Supplementary table 4**. Time divergence calibration and fossil constraint information including comparative literature evidence providing additional support for identified divergence times. First BEAST run divergence estimated with a 25% burn-in whilst second run used a 10%. Sample a1_Ghana is derived from published data [1] | | | | | | |
| --- | --- | --- | --- | --- | --- | --- |
| BEAST Run | Node | Node Calibrated? | Prior Distribution | Constraint Justification | Divergence Estimate (Mean (95% CI)) | Agreement with literature |
| 1^st^ | Culicidae (Outgroup) | Y | *Uniform* min=97 | Poor fossil record and incongruent molecular estimates. Minimum bound at unequivocal *Anophelinae* fossil [2]. | 125.12 (97.00-179.96) | N/A |
| 1^st^ | Brachycera | Y | *Lognormal* mean=3.05, st.dev=0.58, offset=190 | Minimum bound at age of unequivocal brachyceran fossils[3,4]. Peaks 15my prior to incorporate molecular dating work[5–9]. 5% HPD at controversial early fossils[10]. | 202.65 (193.26-214.10) | N/A |
| 1^st^ | Eremoneura | Y | *Lognormal* mean=2.67, st.dev=0.60, offset=164 | Minimum bound at unequivocal eremoneuran fossils[11,12]. Peaks 10my prior to incorporate molecular dating work[5,6]. 5% at earliest credibility date estimated by [5]. | 183.72 (169.48-197.84) | N/A |
| 1^st^ | Cyclorrhapha | Y | *Lognormal* mean=2.73, st.dev=0.65, offset=140 | Minimum bound at unequivocal and taxonomically-debated cyclorrhapan fossils[13,14]. Peaks 10my prior to incorporate molecular dating work[5,6,8]. 5% HPD at earliest credibility date estimated by [5]. | 159.68 (144.27-176.20) | N/A |
| 1^st^ | Schizophora | Y | *Lognormal* mean=3.03, st.dev=0.86, offset=64 | Minimum bound at age of unequivocal schizophoran fossils[15]. Peaks 10my prior to incorporate molecular dating work[5–7,9,16]. 5% HPD at controversial early estimates[8]. | 127.73 (94.74-156.16) | N/A |
| 1^st^ | Drosophila | Y | *Lognormal* mean=1.86, st.dev=0.42, offset=0 | Minimum bound at zero. Peaks 5.4my prior as per the estimate of [17]. 1% HPD at biogeographical constraint for the formation of the *melanogaster* species complex[18,19]. | 12.31 (6.04 – 19.40) | N/A |
| 1^st^ | Stratiomyidae-Tabanidae | N | N/A | N/A | 181.99 (156.47-203.26) | Previous work dates split at ~190mya[5,6] |
| 1^st^ | Stratiomyidae | N | N/A | N/A | 81.07 (45.94-118.84) | Previous work dates split at ~75mya[20,21] |
| 1^st^ | *Hermetia illucens* | N | N/A | N/A | 14.65 (5.46-26.13) | N/A |
| 2^nd^ | Stratiomyidae | Y | *Normal* mean=81.07, st.dev=18.59, offset=0 | Output of BEAST Run 1 | 93.83 (63.17-124.29) | N/A |
| 2^nd^ | Current *Hermetia illucens* root | Y | *Lognormal* mean=2.60, st.dev=0.34, offset=0 | Output of BEAST Run 1 | 6.84 (4.10-9.73) | N/A |
| 2^nd^ | EVE (CAM006568) – a1_Ghana | N | N/A | N/A | 0.02 (0.00 – 0.05) | N/A |
| 2^nd^ | Current lineage *A* root | N | N/A | N/A | 0.59 (0.32 – 0.89) | N/A |
| 2^nd^ | Current lineage *B1* root | N | N/A | N/A | 2.94 (1.80 – 4.15) | N/A |
| 2^nd^ | Africa (CAM006211) – Europe, South (CAM006613) | N | N/A | N/A | 0.60 (0.35 – 0.88) | N/A |
| 2^nd^ | Africa (CAM006211) – Europe, South (CAM006613) – South America (CAM006191) | N | N/A | N/A | 0.70 (0.42 – 1.02) | N/A |
| 2^nd^ | Asia (CAM006206) – Europe, West (CAM006186) | N | N/A | N/A | 0.01 (0.00 - 0.03) | N/A |
| 2^nd^ | Current lineage *B2* root | N | N/A | N/A | 1.22 (0.73 – 1.78) | N/A |
| 2^nd^ | Current lineage *B1* & *B2* (lineage-*b*) root | N | N/A | N/A | 3.79 (2.35 – 5.33) | N/A |

| **Supplementary table 5.** Heterozygosity levels of *Hermetia illucens* populations from around the globe. Samples with sequencing depth >19x and more than one individual per population included. Status allocated based of genomic signatures | | | |
| --- | --- | --- | --- |
| **Population** | **Identified status** | **Sample No. (*n*)** | **Mean heterozygosity** |
| Sheppard: North America | Captive | 4 | 0.032 |
| Sheppard: Europe | Captive | 4 | 0.046 |
| Sheppard: EVE | Captive | 12 | 0.025 |
| North America | Wild | 3 | 0.052 |
| Europe (West) | Wild | 4 | 0.049 |
| Australia | Captive | 4 | 0.046 |
| Africa | Captive | 4 | 0.056 |
| South America | Captive | 4 | 0.087 |

| **Supplementary Table 6.** Number of Runs of Homozygosity identified in the genomes of Black Soldier Fly populations in both captivity and the wild broken down by chromosome. | | | | | | | | | | |
| --- | --- | --- | --- | --- | --- | --- | --- | --- | --- | --- |
| **Chromosome** | **Europe (Sheppard; captive)** | **UK-EVE (captive)** | **North America (Sheppard; captive)** | **Australia (captive)** | **Africa (captive)** | **South America (captive)** | **Asia (wild)** | **Europe (South; wild)** | **Europe (West; wild)** | **North America (wild)** |
| S1 | 26 | 949 | 205 | 63 | 17 | 0 | 2 | 17 | 33 | 4 |
| S2 | 35 | 752 | 117 | 37 | 20 | 2 | 1 | 14 | 45 | 3 |
| S3 | 24 | 622 | 87 | 75 | 26 | 9 | 8 | 13 | 34 | 6 |
| S4 | 17 | 740 | 140 | 50 | 15 | 0 | 0 | 5 | 44 | 0 |
| S5 | 14 | 811 | 89 | 43 | 11 | 1 | 1 | 7 | 34 | 2 |
| S6 | 29 | 460 | 100 | 64 | 16 | 10 | 0 | 21 | 26 | 12 |

| **Supplementary Table 7.** Runs of Homozygosity identified by length (Mb) across the sampled Black Soldier Fly populations in both captivity and the wild. | | | | | | | | | | |
| --- | --- | --- | --- | --- | --- | --- | --- | --- | --- | --- |
| **Length (Mb)** | **Europe (Sheppard; captive)** | **UK-EVE (captive)** | **North America (Sheppard; captive)** | **Australia (captive)** | **Africa (captive)** | **South America (captive)** | **Asia (wild)** | **Europe (South; wild)** | **Europe (West; wild)** | **North America (wild)** |
| 0-0.5 | 132 | 3481 | 612 | 296 | 95 | 22 | 12 | 75 | 194 | 24 |
| 0.5-1 | 12 | 765 | 115 | 32 | 10 | 0 | 0 | 2 | 21 | 2 |
| 1-2 | 1 | 88 | 11 | 3 | 0 | 0 | 0 | 0 | 1 | 1 |
| 2-4 | 0 | 2 | 0 | 1 | 0 | 0 | 0 | 0 | 0 | 0 |

| **Supplementary Table 8.** Summary table for co-ordinates of Runs of Homozygosity (RoH) identified as captive specific hotspots when more than 45% of individuals contained the same RoH region. | | | | | | | |
| --- | --- | --- | --- | --- | --- | --- | --- |
| Chromosome | Start (bp) | End (bp) | Size (bp) | Start (Mb) | End (Mb) | Size (Mp) | Number of Individuals |
| S1 | 193,630,913 | 202,250,373 | 8,619,461 | 193.63 | 202.25 | 8.62 | 20 |
| S1 | 216,573,393 | 219,472,728 | 2,899,336 | 216.57 | 219.47 | 2.90 | 17 |
| S2 | 39,663,035 | 43,055,589 | 3,392,555 | 39.66 | 43.06 | 3.39 | 15 |
| S4 | 16,391,203 | 22,458,359 | 6,067,157 | 16.39 | 22.46 | 6.07 | 17 |
| S5 | 41,766,988 | 48,009,429 | 6,242,442 | 41.77 | 48.01 | 6.24 | 14 |
| S6 | 97,805,609 | 101,468,880 | 3,663,272 | 97.81 | 101.47 | 3.66 | 16 |
| S6 | 97,104,344 | 97,805,464 | 701,121 | 97.10 | 97.81 | 0.70 | 15 |

| **Supplementary table 9.** Genomic regions of introgression from domesticated populations into wild European (South) from the North America Sheppard breed, using Africa as P1. Genes (*n*=85) found within the 2.1 Mb introgressed region (S1: 173,322,084 - 175,418,642) are identified with predicted functions. | | | |
| --- | --- | --- | --- |
| **Gene ID** | **Copies (#)** | **Gene name** | **Function** |
| *phm* | *15* | cytochrome P450 6A1-like (*phantom*) | Required for cuticle secretion, head involution and dorsal closure. |
| *Cyp6a2* | 2 | cytochrome P450 6a2-like (*Cyp6a2*) | Involved in the breakdown of exogenous compounds such as insecticides and insect hormones. May include DDT and caffeine breakdown. |
| *Cyp6a8* | 8 | cytochrome P450 6a8-like | Involved in the breakdown of exogenous compounds such as insecticides and insect hormones. Including lauric acid and caffeine. |
| *Cyp6a9* | 6 | cytochrome P450 6a9-like | Involved in the breakdown of exogenous compounds such as insecticides and insect hormones. |
| *Cyp6a13* | 7 | probable cytochrome P450 6a13 | Involved in the breakdown of exogenous compounds such as insecticides and insect hormones. Also involved in defence responses against bacteria. |
| *Cyp6a14* | 19 | probable cytochrome P450 6a14 | Involved in the breakdown of exogenous compounds such as insecticides and insect hormones. |
| *Cyp6a18* | 1 | probable cytochrome P450 6a18 | Involved in the breakdown of exogenous compounds such as insecticides and insect hormones. |
| *Cyp6a20* | 8 | probable cytochrome P450 6a20 | Involved in the breakdown of aggression promoting pheromones. |
| *Cyp6a21* | 10 | probable cytochrome P450 6a21 | Involved in the breakdown of exogenous compounds such as insecticides and insect hormones. |
| *Cyp6d4* | 1 | probable cytochrome P450 6d4 | Involved in the breakdown of exogenous compounds such as insecticides and insect hormones. |
| *Cyp6d5* | 2 | probable cytochrome P450 6d5 | Involved in the breakdown of exogenous compounds such as insecticides and insect hormones. |
| *bowel* | 1 | Brother of odd with entrails limited (*bowel*) | Required for embryonic fore & hindgut morphogenesis in the embryo. |
| *dac* | 1 | dachshund (*dac*) | Regulates eye, leg, gonad and brain development. |
| *AaSDR-1* | 1 | farnesol dehydrogenase-like | Oxidates farnesol into farnesal, a JH precursor. |
| *Cdep* | 1 | FERM, ARHGEF and pleckstrin domain-containing protein 2 (*Cdep*) | Predicted to enable guanyl-nucleotide exchange factor activity |
| *Atf6* | 1 | cyclic AMP-dependent transcription factor ATF-6 alpha | Involved in endoplasmic reticulum unfolded protein response and regulation of transcription by RNA polymerase II |
| *NA* | 1 | unnamed | NA |

| **Supplementary table 10.** Genes within the introgressed region across chromosome three from a wild Asian representative into the domesticated industry-strain-A. | |
| --- | --- |
| **Gene** | **Genes names of interest** |
| jg10537.t1 | unnamed |
| jg10538.t1 | coiled-coil domain-containing protein 43 |
| jg10539.t1 | nucleolar transcription factor 1 |
| jg10540.t1 | ATP-binding cassette sub-family C member Sur |
| jg10541.t1 | cGMP-dependent protein kinase 1 isoform X1 |
| jg10542.t1 | uncharacterized |
| jg10543.t1 | uncharacterized |
| jg10544.t1 | unnamed |
| jg10545.t1 | N-acetylneuraminate lyase B-like |
| jg10546.t1 | protein obstructor-E |
| jg10547.t1 | pickpocket protein 28 isoform X1 |
| jg10548.t1 | trifunctional purine biosynthetic protein adenosine-3 |
| jg10549.t1 | dnaJ homolog subfamily C member 28 |
| jg10550.t1 | translocation protein SEC62 isoform X2 |
| jg10551.t1 | importin subunit alpha |
| jg10552.t1 | ovarian-specific serine/threonine-protein kinase Lok isoform X2 |
| jg10553.t1 | malate dehydrogenase, mitochondrial-like |
| jg10554.t1 | interferon-inducible double-stranded RNA-dependent protein kinase activator A homolog isoform X1 |
| jg10555.t1 | SUZ domain-containing protein 1-like |
| jg10556.t1 | valois |
| jg10557.t1 | glycine--tRNA ligase |
| jg10558.t1 | uncharacterized |
| jg10559.t1 | dual oxidase maturation factor 1 |
| jg10560.t1 | dual oxidase |
| jg10561.t1 | glycoprotein-N-acetylgalactosamine 3-beta-galactosyltransferase 1-like isoform X1 |
| jg10562.t1 | NA |
| jg10563.t1 | glycoprotein-N-acetylgalactosamine 3-beta-galactosyltransferase 1-like isoform X1 |
| jg10564.t1 | dynactin subunit 5 |
| jg10565.t1 | probable phosphorylase b kinase regulatory subunit beta isoform X4 |
| jg10566.t1 | short-chain specific acyl-CoA dehydrogenase, mitochondrial-like |
| jg10567.t1 | transient receptor potential-gamma protein isoform X4 |
| jg10568.t1 | THO complex subunit 2 isoform X1 |
| jg10569.t1 | THO complex subunit 2 isoform X3 |
| jg10570.t1 | sarcoplasmic calcium-binding protein, alpha-B and -A chains |
| jg10571.t1 | uncharacterized |
| jg10572.t1 | uncharacterized |
| jg10573.t1 | NPC intracellular cholesterol transporter 1 isoform X2 |
| jg10574.t1 | piezo-type mechanosensitive ion channel component isoform X5 |
| jg10575.t1 | drosulfakinins |
| jg10576.t1 | facilitated trehalose transporter Tret1-like isoform X2 |
| jg10577.t1 | uncharacterized |
| jg10578.t1 | unnamed |
| jg10579.t1 | uncharacterized |
| jg10580.t1 | uncharacterized |
| jg10581.t1 | uncharacterized |
| jg10582.t1 | enhancer of mRNA-decapping protein 4 homolog isoform X4 |
| jg10583.t1 | unnamed |
| jg10584.t1 | opsin-3-like |
| jg10585.t1 | NA |
| jg10586.t1 | piggyBac transposable element-derived protein 4-like |
| jg10587.t1 | leucine-rich repeat extensin-like protein 3 |
| jg10588.t1 | serine/threonine-protein kinase meng-po |
| jg10589.t1 | sphingolipid delta(4)-desaturase DES1 |
| jg10590.t1 | solute carrier family 12 member 9 |
| jg10591.t1 | solute carrier organic anion transporter family member 74D |
| jg10592.t1 | solute carrier organic anion transporter family member 74D-like |
| jg10593.t1 | eukaryotic peptide chain release factor GTP-binding subunit ERF3A-like isoform X1 |
| jg10594.t1 | eukaryotic peptide chain release factor GTP-binding subunit ERF3A-like isoform X3 |
| jg10595.t1 | eukaryotic peptide chain release factor GTP-binding subunit ERF3A-like isoform X2 |
| jg10596.t1 | DNA fragmentation factor subunit beta |
| jg10597.t1 | probable phospholipid-transporting ATPase IIA isoform X3 |
| jg10598.t1 | 39S ribosomal protein L55 |
| jg10599.t1 | transmembrane protease serine 9-like |
| jg10600.t1 | outer dense fiber protein 2 |
| jg10601.t1 | transmembrane protease serine 9 |
| jg10602.t1 | putative beta-carotene-binding protein |
| jg10603.t1 | ankyrin repeat domain-containing protein 49 |
| jg10604.t1 | uncharacterized |
| jg10605.t1 | kinesin-like protein KIF21A isoform X1 |
| jg10606.t1 | unnamed |
| jg10607.t1 | collagen alpha-3(IX) chain-like isoform X2 |
| jg10608.t1 | NA |
| jg10609.t1 | collagen alpha-2(IX) chain-like |
| jg10610.t1 | unnamed |
| jg10611.t1 | collagen alpha-1(X) chain-like |
| jg10612.t1 | collagen alpha-1(I) chain-like |
| jg10613.t1 | unnamed |
| jg10614.t1 | unnamed |
| jg10615.t1 | irregular chiasm C-roughest protein |
| jg10616.t1 | uncharacterized |
| jg10617.t1 | probable ATP-dependent RNA helicase DDX23 |
| jg10618.t1 | zinc finger matrin-type protein 5 |
| jg10619.t1 | basic helix-loop-helix transcription factor amos |
| jg10620.t1 | unnamed |
| jg10621.t1 | FAST kinase domain-containing protein 1 |
| jg10622.t1 | NA |
| jg10623.t1 | unnamed |
| jg10624.t1 | unnamed |
| jg10625.t1 | protein RCC2 homolog |
| jg10626.t1 | D(3) dopamine receptor |
| jg10627.t1 | unnamed |
| jg10628.t1 | probable salivary secreted peptide |
| jg10629.t1 | probable salivary secreted peptide |
| jg10630.t1 | probable salivary secreted peptide |
| jg10631.t1 | probable salivary secreted peptide |
| jg10632.t1 | probable salivary secreted peptide |
| jg10633.t1 | microtubule-associated protein futsch-like |
| jg10634.t1 | FAD-dependent oxidoreductase domain-containing protein 1-like |
| jg10635.t1 | FAD-dependent oxidoreductase domain-containing protein 1-like |
| jg10636.t1 | probable 26S proteasome non-ATPase regulatory subunit 3 |
| jg10637.t1 | unnamed |
| jg10638.t1 | myb-like protein Q |
| jg10639.t1 | dynein assembly factor 6, axonemal |
| jg10640.t1 | cysteine-rich venom protein-like isoform X3 |
| jg10641.t1 | protein phosphatase PHLPP-like protein |
| jg10642.t1 | LIM/homeobox protein Lhx3 isoform X4 |
| jg10643.t1 | NA |
| jg10644.t1 | protein anon-37Cs |
| jg10645.t1 | NA |
| jg10646.t1 | pre-mRNA-splicing factor ATP-dependent RNA helicase DHX16 |
| jg10647.t1 | protein l (2)37Cc |
| jg10648.t1 | general transcription factor 3C polypeptide 5 |
| jg10649.t1 | NA |
| jg10650.t1 | adenylate kinase isoenzyme 1 isoform X2 |
| jg10651.t1 | lysosome-associated membrane glycoprotein 1 |
| jg10652.t1 | unnamed |
| jg10653.t1 | unnamed |
| jg10654.t1 | RNA-binding protein 5-B-like isoform X1 |
| jg10655.t1 | cysteine-rich protein 2-binding protein |
| jg10656.t1 | dnaJ protein homolog 1 |
| jg10657.t1 | myosin-4-like |

| **Supplementary table 11**. Diptera species used in the initial BEAST dating run. Also shown is the dipteran clade within which each species falls, and thus which calibrated nodes they feature in (noting the nested nature of the dipteran phylogeny means that Drosophila individuals fall within Schizophora, which falls within Cyclorrhapha, which falls within Eremoneura, which falls within Brachycera). The GenBank accession numbers for publicly available mitogenomes are given expect sample a1_Ghana from [1]. | | | | |
| --- | --- | --- | --- | --- |
| BEAST Run | Species | BSF mt-lineage | Node | GenBank accession number |
| 1 | *Aedes aegypti* | N/A | Culicidae (Outgroup) | NC_035159.1 |
| 1 | *Anopheles gambiae* | N/A | Culicidae (Outgroup) | NC_035159.1 |
| 1 | *Cydistomyia duplonotata* | N/A | Brachycera | NC_008756.1 |
| 1 | *Hermetia illucens* (EVE8) | Lineage *A* | Brachycera; Stratiomyidae; Hermetia illucens | This study |
| 1 | *Hermetia illucens* (AF1) | Lineage *B2* | Brachycera; Stratiomyidae; Hermetia illucens | This study |
| 1 | *Ptecticus aurifer* | N/A | Brachycera; Stratiomyidae | This study |
| 1 | *Hercostomus brevipilosus* | N/A | Brachycera; Eremoneura | NC_046942.1 |
| 1 | *Megaselia scalaris* | N/A | Brachycera; Eremoneura; Cyclorrhapha | NC_023794.1 |
| 1 | *Dermatobia hominis* | N/A | Brachycera; Eremoneura; Cyclorrhapha; Schizophora | NC_006378.1 |
| 1 | *Melophagus ovinus* | N/A | Brachycera; Eremoneura; Cyclorrhapha; Schizophora | NC_037368.1 |
| 1 | *Musca domestica* | N/A | Brachycera; Eremoneura; Cyclorrhapha; Schizophora | NC_024855.1 |
| 1 | *Drosophila melanogaster* | N/A | Brachycera; Eremoneura; Cyclorrhapha; Schizophora; *Drosophila* | NC_024511.2 |
| 1 | *Drosophila simulans* | N/A | Brachycera; Eremoneura; Cyclorrhapha; Schizophora; *Drosophila* | NC_005781.1 |
| 2 | *Ptecticus aurifer* | N/A | Brachycera; Stratiomyidae | This study |
| 2 | *Hermetia illucens* (CAM006568) | Lineage *A* | Brachycera; Stratiomyidae; *Hermetia illucens* | This study |
| 2 | *Hermetia illucens* (a1_Ghana) | Lineage *A* | Brachycera; Stratiomyidae; *Hermetia illucens* | Not supplied [1] |
| 2 | *Hermetia illucens* (CAM006196) | Lineage *A* | Brachycera; Stratiomyidae; *Hermetia illucens* | This study |
| 2 | *Hermetia illucens* (CAM006202) | Lineage *B1* | Brachycera; Stratiomyidae; *Hermetia illucens* | This study |
| 2 | *Hermetia illucens* (CAM006181) | Lineage *B1* | Brachycera; Stratiomyidae; *Hermetia illucens* | This study |
| 2 | *Hermetia illucens* (CAM006206) | Lineage *B2* | Brachycera; Stratiomyidae; *Hermetia illucens* | This study |
| 2 | *Hermetia illucens* (CAM006211) | Lineage *B2* | Brachycera; Stratiomyidae; *Hermetia illucens* | This study |
| 2 | *Hermetia illucens* (CAM006186) | Lineage *B2* | Brachycera; Stratiomyidae; *Hermetia illucens* | This study |
| 2 | *Hermetia illucens* (CAM006613) | Lineage *B2* | Brachycera; Stratiomyidae; *Hermetia illucens* | This study |
| 2 | *Hermetia illucens* (CAM006191) | Lineage *B2* | Brachycera; Stratiomyidae; *Hermetia illucens* | This study |
